# Genome Assembly and Population Resequencing Reveal the Geographical Divergence of ‘Shanmei’ (*Rubus corchorifolius*)

**DOI:** 10.1101/2021.11.22.469527

**Authors:** Yinqing Yang, Kang Zhang, Ya Xiao, Lingkui Zhang, Yile Huang, Xing Li, Shumin Chen, Yansong Peng, Shuhua Yang, Yongbo Liu, Feng Cheng

## Abstract

***Rubus corchorifolius*** (‘**Shanmei**’ or mountain berry, 2n =14) is widely distributed in China, and its fruit has high nutritional and medicinal values. Here, we report a high-quality chromosome-scale **genome assembly** of Shanmei, with a size of 215.69 Mb and encompassing 26,696 genes. Genome comparisons among Rosaceae species show that Shanmei and Fupenzi (*Rubus chingii* Hu) are most closely related, and then is blackberry (*Rubus occidentalis*). Further **resequencing** of 101 samples of Shanmei collected from four regions in provinces of Yunnan, Hunan, Jiangxi, and Sichuan in South China reveals that the Hunan population of Shanmei possesses the highest diversity and may represent the relatively more ancestral population. Moreover, the Yunnan population undergoes strong selection based on nucleotide diversity, linkage disequilibrium, and the historical effective population size analyses. Furthermore, genes from candidate genomic regions that show strong **divergence** are significantly enriched in flavonoid biosynthesis and plant hormone signal transduction, indicating the genetic basis of adaptation of Shanmei to the local environments. The high-quality genome sequences and the variome dataset of Shanmei provide valuable resources for breeding applications and for elucidating the **genome evolution** and ecological adaptation of *Rubus* species.

## Introduction

*Rubus corchorifolius*, also named ‘Shanmei’, belongs to the Rosaceae family. *Rubus* is a large genus consisting of approximately 750 species, most of which are perennial shrubs and biennial vines [1]. Species from *Rubus* constitute important components of the ground layer of hillsides, valleys, and large forest canopy gaps, providing a host of ecological benefits, including soil stabilization, reduced soil nutrient loss, as well as food for wildlife. The wide distribution of *Rubus* species is accompanied by rich diversity both in terms of stress adaptation and organ development, and thus *Rubus* has great potential for agricultural utilization [2]. There are 201 species and 98 varieties of *Rubus* distributed in various regions of China, which provide important resources for the exploration of the biological diversity of the genus. At present, only a few species in the genus *Rubus*, including blackberry, dewberry, and arctic raspberry, have been domesticated and utilized in breeding programs [3]. Some of them, such as blackberry, have been developed as important crops with great economic value [4].

Shanmei, one of the most important *Rubus* species with many desirable horticultural traits, is widely distributed in China. There are rich diversities and significant differences among Shanmei population from different geographic regions, including in characters associated with environmental adaptation, population size, as well as flowering, single fruit weight, and fruit size. The fruit of Shanmei is popular for its unique flavor and nutrients, such as high amounts of anthocyanins, superoxide dismutase (SOD), vitamin C, and essential amino acids [5, 6]. Shanmei fruit has been processed into food products as jam, juice, wine, and ice cream, and is becoming increasingly popular among consumers [7]. The terpenoids extracted from Shanmei leaves can suppress the development of cancer cells by inducing tumor cell differentiation and apoptosis [8]. Considering the important economic and medicinal values of Shanmei, it is of practical significance to explore and utilize its wild resources.

The genome is an essential resource for studying the traits and gene functions of species [9]. Thus far, several high-quality genomes of Rosaceae species have been released, including *Pyrus communis* (pear) [10], *Malus domestica* (apple) [11], *Prunus persica* (peach) [11], *Prunus mume* (plum) [12], *Prunus armeniaca* (apricot) [13], *Fragaria vesca* (strawberry) [14], *Rosa chinensis* (rose) [15], *Rubus occidentalis* (blackberry) [4], and *Rubus chingii* Hu (Fupenzi) [16]. Comparative genomics analysis revealed the evolutionary relationships among Rosaceae species and reconstructed a hypothetical ancestry. Using the data of chloroplast genome, previous work showed that Shanmei was located at the Rubeae clade of the Rosaceae family, and is closest to *Rubus rufus* [17]. Genomic data also provide important genetic resources for the identification of important agronomic traits including flavor, scent, nutritional value, flower color, and flowering times [10, 13, 14]. Blackberry and Fupenzi, which are closely related to Shanmei, are the two members of *Rubus* with a chromosome-level genome [4, 16]. The gene duplication of chalcone synthase (*CHS*), the first committed enzyme in flavonoid biosynthesis, was found to be positively correlated with trait domestication in blackberry based on genomic resources [4]. In addition, transcriptome data were used to analyze gene expression patterns during blackberry fruit ripening. These findings contributed to our understanding of the biology and breeding application of blackberry [4]. For Fupenzi, the genome analysis revealed that there was a tandem gene cluster in chromosome 02 that regulated the biosynthetic pathway of hydrolyzable tannins [16]. However, as there are no available genomic resources for Shanmei, the investigations of the genetic mechanisms underlying the favorable traits or the exploitations on population resource of this potential species as a fast-growing economical horticultural crop is hindered.

In this study, we generated the first chromosome-scale assembly of the Shanmei genome and re-sequenced 101 Shanmei samples collected from four different geographical regions in China. Comparative genomics analysis revealed the expanded gene families that allow Shanmei to occupy its special ecological niche. The population analysis found that the Hunan population is the relatively ancestral group, while the Yunnan population underwent strong selection. The high-quality genome and population variome dataset of Shanmei not only provided insights into its evolution and geographical divergence but also provided a foundation for the breeding utilization of Shanmei.

## Results

### Pseudo-chromosome construction of the Shanmei genome

We sequenced and assembled the genome of Shanmei (**Figure 1A**) using combined sequencing data from Oxford Nanopore Technologies (ONT), Illumina HiSeq, and high-throughput chromosome conformation capture (Hi-C). The genome was estimated to be 187.82 Mb in size with a heterozygosity ratio of 1.82% based on 21-kmer counting, showing that it is highly heterozygous (Figure S1). A total of 36.87 Gb (∼180 **×**) Nanopore long reads were generated and assembled into 221 contigs (Table S1). The size of the assembly was 330 Mb, with a contig N50 of 2.49 Mb. It was speculated that the significantly larger size of the assembly compared to the estimate was caused by the introduction of the heterozygous contigs, considering the high heterozygosity ratio. Therefore, redundant contigs were then identified and filtered out using Purge Haplotigs (version 1.2.3) [18], and only 120 contigs (215.69 Mb) were retained for further analysis. A total of 43.56 Gb (∼220 ×) Hi-C data were further used to link the contigs into scaffolds. Consequently, 10 scaffolds were obtained with an N50 of 29.50 Mb. The seven largest scaffolds comprised 117 contigs, which accounted for 99.35% (214.29 Mb) of the assembled genome and were corresponding to the seven pseudo-chromosomes of Shanmei (Figure 1B, Figure S2; Table S2). Furthermore, the telomere sequences were identified in the ends of the seven chromosomes (Figure S2), which supported the high quality of the genome assembly of Shanmei. Additionally, Benchmarking Universal Single-Copy Ortholog (BUSCO) analysis showed that 94.7% of the BUSCO genes were successfully identified in the Shanmei genome (Table 1).

**Table 1.** Assembly and annotation statistics of the Shanmei genome

| Type | Contig |  | Scaffold |  |
| --- | --- | --- | --- | --- |
|  | Size (Mb) | Number | Size (Mb) | Number |
| Maximum | 11.08 | 1 | 36.68 | 1 |
| N50 | 3.34 | 21 | 29.50 | 4 |
| N90 | 0.78 | 80 | 27.00 | 6 |
| Total length | 215.69 | 120 | 215.74 | 10 |
| Chromosomes | / | / | 214.29 (99.35%) |  |
| Genes | / | / | / | 26,696 |
| Transposable<br>elements | / | / | 77.33 (35.85%) | / |

**Figure 1.**
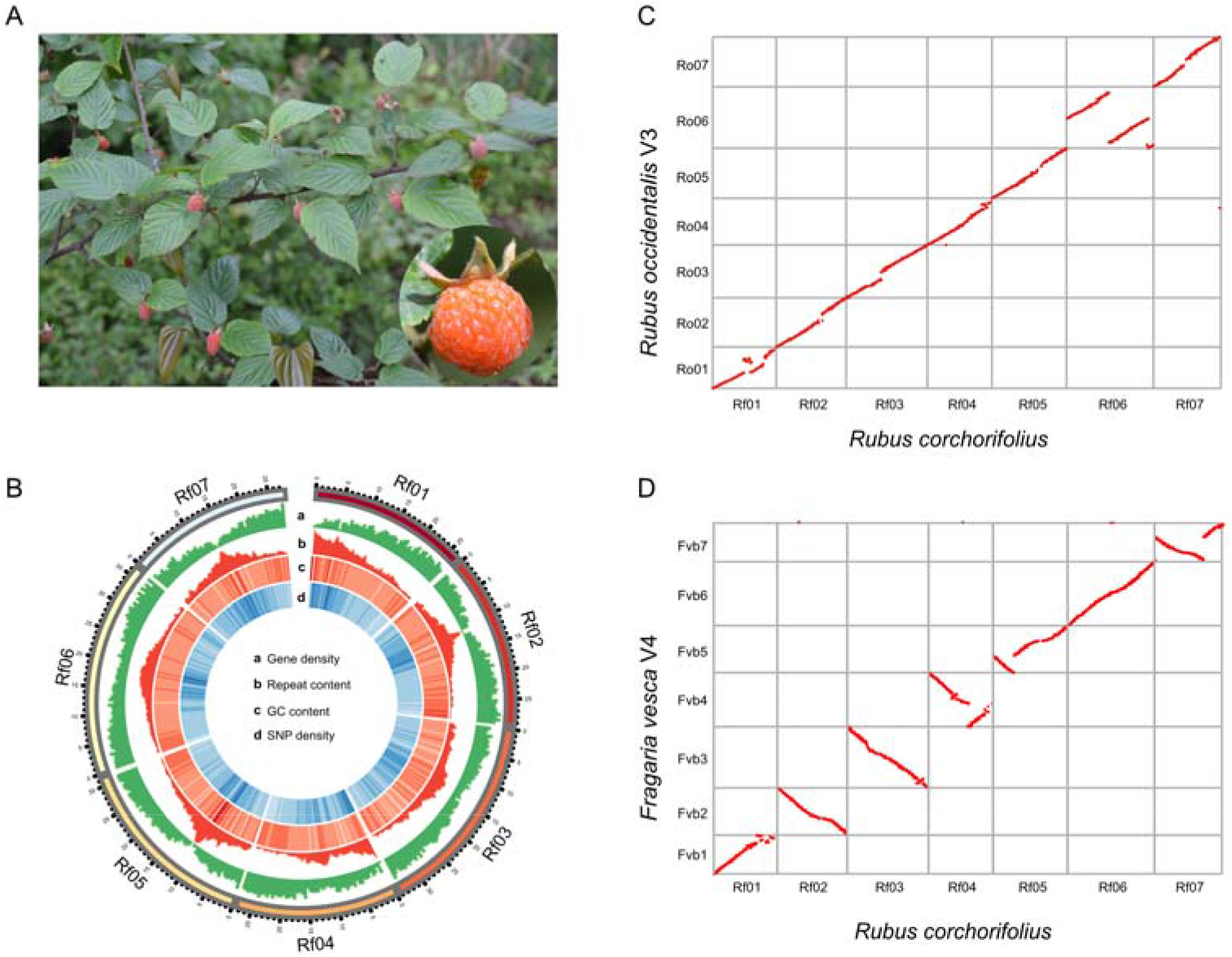
Assembly and characterization of the Shanmei genome. **A**. The Shanmei plant and the close-up view of its fruit. **B**. The landscape of the Shanmei genome. a: gene density; b: repeat content; c: GC content; and d: SNP density. The chromosome units are in 1 Mb. **C**. Genomic synteny between Shanmei and blackberry. **D**. Genomic synteny between Shanmei and strawberry. *Rubus corchorifolius*, Shanmei; *Rubus occidentalis*, blackberry; *Fragaria vesca*, strawberry.

We employed an integrated pipeline to annotate the genome by combining *de novo* prediction, homology search, and RNA-seq data alignment (Methods). A total of 26,696 protein-coding genes were predicted in the Shanmei genome (Table S3). The high gene prediction quality was supported by the fact that 1976 (93.1%) of the BUSCO genes were found in the Shanmei gene set. In addition, repeat annotation revealed that approximately 35.85% (77.33 Mb) of the genome was composed of repetitive elements, comparable to that of Fupenzi [16]. The predominant type of transposable elements (TEs) was long terminal repeat (LTR) retrotransposons, accounting for 11.26% of the genome (Table S4).

### Expanded gene families in flavonoid biosynthesis and stress resistance

Rosaceae is an economically important family composed of 2800 species among 95 genera, including the specialty fruit crops apple, almond, and blackberry. In order to infer the phylogenetic position of Shanmei in Rosaceae, we obtained 932 single-copy genes and constructed a phylogenetic tree for 10 Rosaceae species with grape as the outgroup (**Figure 2A**). The tree showed that Shanmei and Fupenzi were most closely related. Each genomic region in Shanmei was found to be orthologous to a single region in each of Fupenzi, blackberry, and strawberry based on genomic synteny analysis, suggesting that no lineage-specific genome duplication occurred in Shanmei after the common γ hexaploidization event [4] (Figure 1C and D, Figure S3 and 5). In addition, a large translocation on chromosome 6 between Shanmei and blackberry was identified (Figure 1C). To verify the accuracy of the assembly results, we re-adjusted the scaffolding orders of the contigs in chromosome 6 to follow those in blackberry and found that the resultant Hi-C heatmap exhibited clear mis-connections (Figure S4), suggesting an authentic translocation between Shanmei and blackberry, which may be associated with the divergence of the two species.Meanwhile, four smaller inversions on chromosome 1 and 4 were found between Shanmei and Fupenzi (Figure S5). These inversions were also verified based on Hi-C heatmap (Figure S6 and 7).

**Figure 2.**
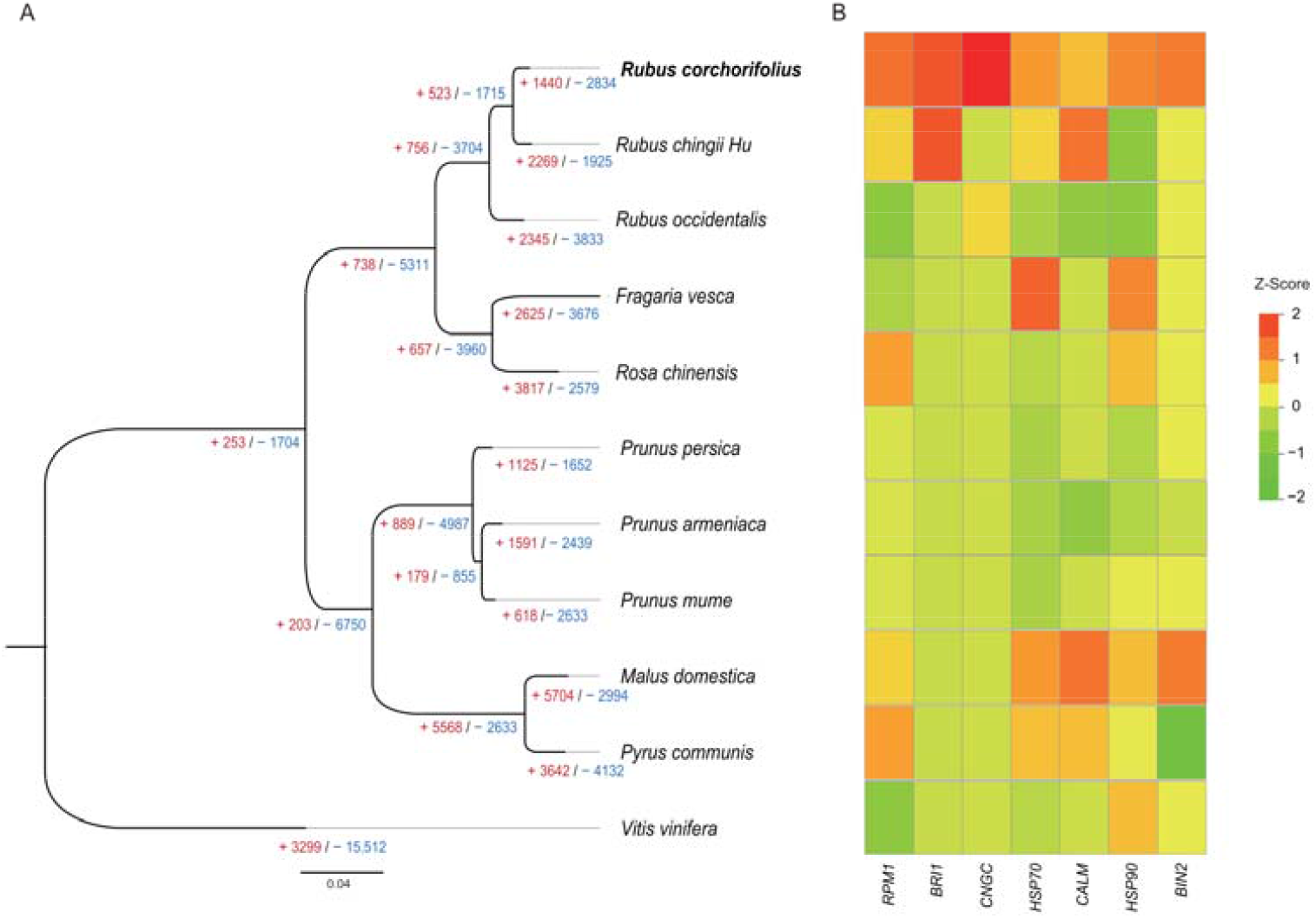
Phylogenetic position and gene family expansion of Shanmei. **A**. The phylogenetic tree of Shanmei and eight other Rosaceae species built based on 897 single-copy genes, with *Vitis vinifera* as the outgroup. The inferred expansion (red numbers) and contraction (blue numbers) of gene families in different genomes are indicated. **B**. Copy number variations of the gene family associated with environmental adaptation.

Subsequently, we determined the expansion and contraction of orthologous gene families using CAFÉ (version 4.2.1) [19] software. It was found that a total of 1440 and 2834 gene families underwent expansion and contraction, respectively in Shanmei. Kyoto Encyclopedia of Genes and Genomes (KEGG) enrichment analyses revealed that the expanded gene families mainly participated in phenylalanine metabolism, flavonoid biosynthesis, brassinosteroid biosynthesis, and biosynthesis of secondary metabolites (corrected *P*-value□<□0.05; Figure S8A; Table S5). In contrast, the contracted gene families were mainly associated with monoterpenoid biosynthesis, alpha-linolenic acid metabolism, and nitrogen metabolism (corrected *P*-value□<□0.05; Figure S8B; Table S6). Furthermore, some significantly expanded genes were closely related to stress resistance, such as *HSP90, HSP70, BRI1, BIN2*, and *RPM1* (Figure 2B; Table S7). Multiple studies have reported that the expanded families in different plants may contribute to abiotic and biotic stress tolerance. Overexpression of *OsHsp90* can enhance cell viability and heat tolerance in rice under heat stress [20]. BRI1, as a signal receptor in the brassinosteroid signal transduction pathway, plays an important role in plant development and disease resistance [21]. *RPM1* is a resistance gene that improves the resistance to root-knot nematodes in wild *myrobalan plum* [22]. In summary, the expansion of these genes may contribute to the environmental adaptability of Shanmei in the wild.

### Genomic variation response to morphology

Shanmei is a low shrub, which is typical in the genus *Rubus*. Lignin is an important factor associated with plant height differences [23]. We identified the key genes for lignin biosynthesis in Shanmei, based on the homologous genes reported in *Arabidopsis thaliana* (Arabidopsis). We found that the gene copy numbers of *CAD* (*P* value: 0.036) and *COMT* (*P* value: 0.047) were increased significantly in trees (Figure S9; Table S9). There are nine copies of *CAD* in Shanmei and 12 in strawberry, comparing to 24 and 18 in trees of pear and peach, respectively. It is known that the decreased expression dosage of *CAD* leads to sterility and dwarfing in Arabidopsis [24]. The increased copy number of *CAD* genes in trees may contribute to their activity of lignin biosynthesis. Meanwhile, the number of *COMT* in shrubs (eight in Shanmei) was more than that in herbs (five in strawberry), and both were less than that in trees (15 in pear; 14 in peach) (Figure S9C). *COMT* is one of the important enzymes controlling lignin monomer production in plant cell wall biosynthesis, and decreased expression of *COMT* resulted in decreased lignin content [25]. Furthermore, we compared the expression of genes related to lignin biosynthesis in the stem organ of three representative species, i.e., strawberry, Shanmei, and pear, and found that the expression level of lignin biosynthesis-related genes showed a positive association with the heights of species that were compared (Figure S9D), which further supported the dosage effect of these lignin biosynthesis-related genes in Rosaceae.

Anthocyanins are abundant in Shanmei and have essential functions in stress resistance and fruit coloring. We identified the key genes for anthocyanin biosynthesis in Shanmei genome based on anthocyanin-related gene pathways reported in Arabidopsis and blackberry [26, 27] (Methods; Table S8). Among them, *MYB10* is the main regulator in anthocyanin biosynthesis. By comparing the functional domain of MYB10 from 10 species of Rosaceae, we identified two conservative motifs (R2 and R3 as shown in Figure S10A) in MYB10, and found that the amino acids of alanine (A) in R3 motif was substituted by serine (S) in Shanmei, which is shared only by red raspberry and blackberry [27]. In addition, we found a novel substitution in which the aspartic acid (D, acidic amino acid) located in the R3 motif was replaced by the asparagine (N, neutral amino acid) only in blackberry (Figure S10B).

### Population structure of Shanmei

To elucidate the population structure of Shanmei, we collected 101 samples from the provinces of Jiangxi (21), Hunan (25), Yunnan (25), and Sichuan (30) in South China (Methods), which is the main distribution area of Shanmei. We resequenced these samples at an average depth of 34-fold coverage (**Figure 3A**; Table S10). The resultant average mapping rate was 91.7% (Table S10). Single nucleotide polymorphisms (SNPs) were identified with the Genome Analysis Toolkit (GATK) [28]. After filtering, a total of 758,978 SNPs were retained for further analysis. The SNPs were evenly distributed across the chromosomes (Figure 1d; Table S11). A total of 18.98% and 10.56% of the SNPs were located in gene-proximal (2 kb upstream or downstream of a coding sequence) and in coding regions, respectively. Moreover, a total of 38,468 (5.07%) SNPs resulted in non-synonymous sequence changes, among which 837 (0.11%) SNPs disrupted the coding sequence (premature stop codon).

**Figure 3.**
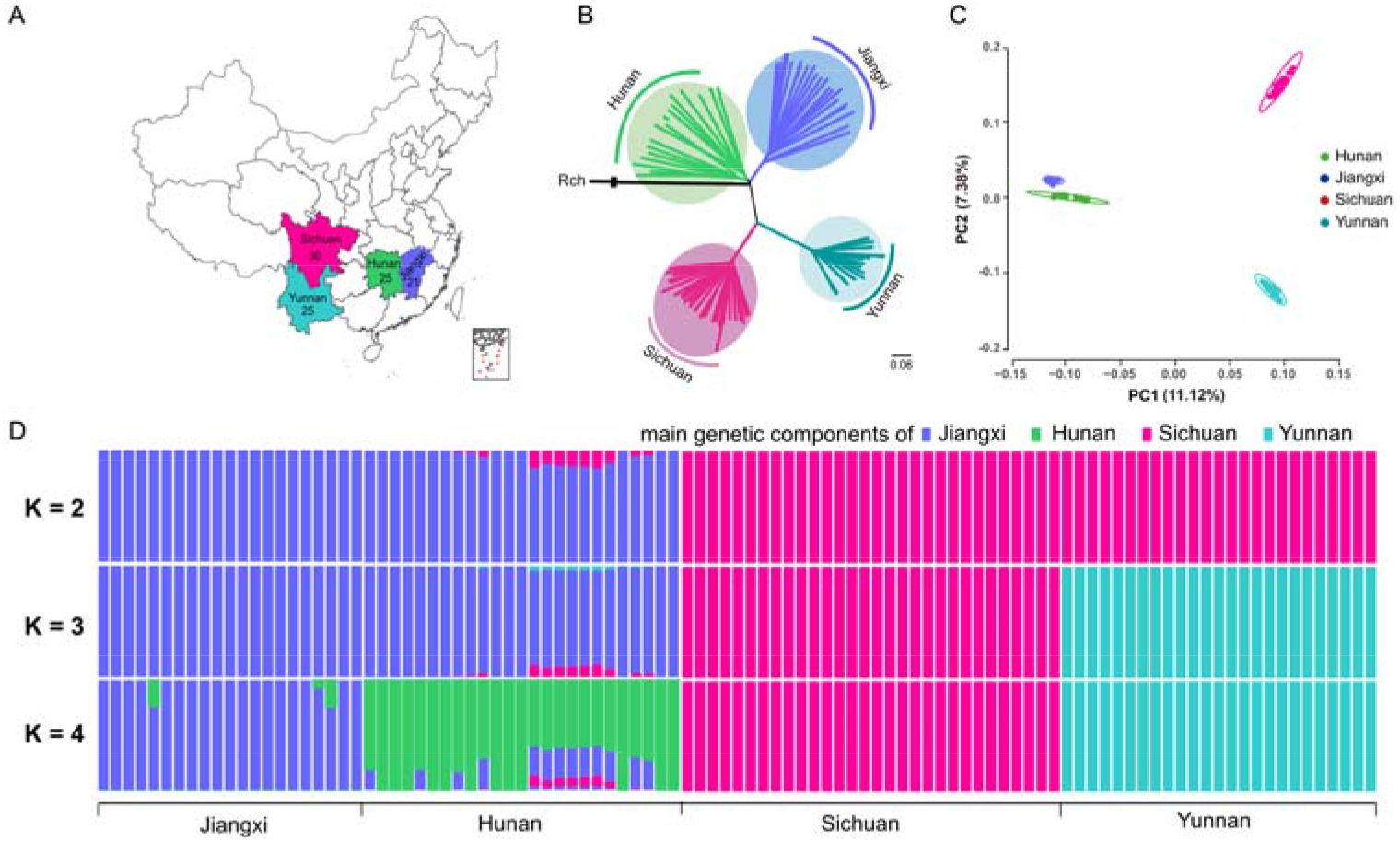
Population structure of Shanmei. **A**. The geographic locations sampled in this study. The numbers denote the number of samples collected in the corresponding region. **B**. Best maximum-likelihood tree showing the phylogenetic relationships of the 101 Shanmei samples. The genome of Fupenzi (Rch) was used as the outgroup. **C**. Principal component analysis of the Shanmei populations. PC1 and PC2 split populations into four clusters. **D**. Genetic admixture of the Shanmei samples analyzed. The length of each colored segment represents the proportion of genetic components in each sample (K = 2–4). Blue, green, pink, and cyan represent the main genetic components of Jiangxi, Hunan, Sichuan, and Yunnan Shanmei groups, respectively.

In order to further explore the phylogenetic relationships among the 101 samples, we constructed a phylogenetic tree based on the maximum-likelihood (ML) method and found that the accessions were clustered into four clades, which exactly corresponded to four geographical regions (Figure 3B). Principal component analysis (PCA) also revealed four clusters, which was consistent with the phylogenetic result (Figure 3C). We found that the Jiangxi and Hunan groups remained closely associated. The genetic clustering results were further confirmed by the genetic structure analyses (Figure 3D). When K= 4, the same four groups were observed, indicating the distinguishable divergence among populations from different geographical regions. These data showed that the Hunan population is more diversified and may represent the relatively ancestral group of Shanmei.

### Flavonoid and phytohormone pathways contributed to the adaptation of Shanmei

On the basis of the phylogenetic relationships and population structure, we further investigated the population-level heterozygosity among Shanmei populations. We found that the Yunnan group had a lower level of heterozygosity than groups of Jiangxi, Sichuan, and Hunan (**Figure 4A**; Table S12). Consistently, the linkage disequilibrium (LD, indicated by *r*^2^) decay rate was highest in the Yunnan group followed by the Sichuan, Jiangxi, and Hunan groups (Figure 4B). We then calculated the nucleotide diversities (π) for the four groups. The Yunnan group had the lowest nucleotide diversity (π = 6.0 × 10^−4^) compared with groups of Sichuan (π = 7.5 × 10^−4^), Hunan (π = 8.1 × 10^−4^), and Jiangxi (π = 8.5 × 10^−4^) (Figure 4C). In addition, the historical effective population size analysis showed that the population size of Yunnan decreased significantly in the recent period compared to the other groups (Figure 4D). In short, these results suggested that the Yunnan group, which is distributed at the high altitude region, underwent the greatest pressure of selection among the four groups.

**Figure 4.**
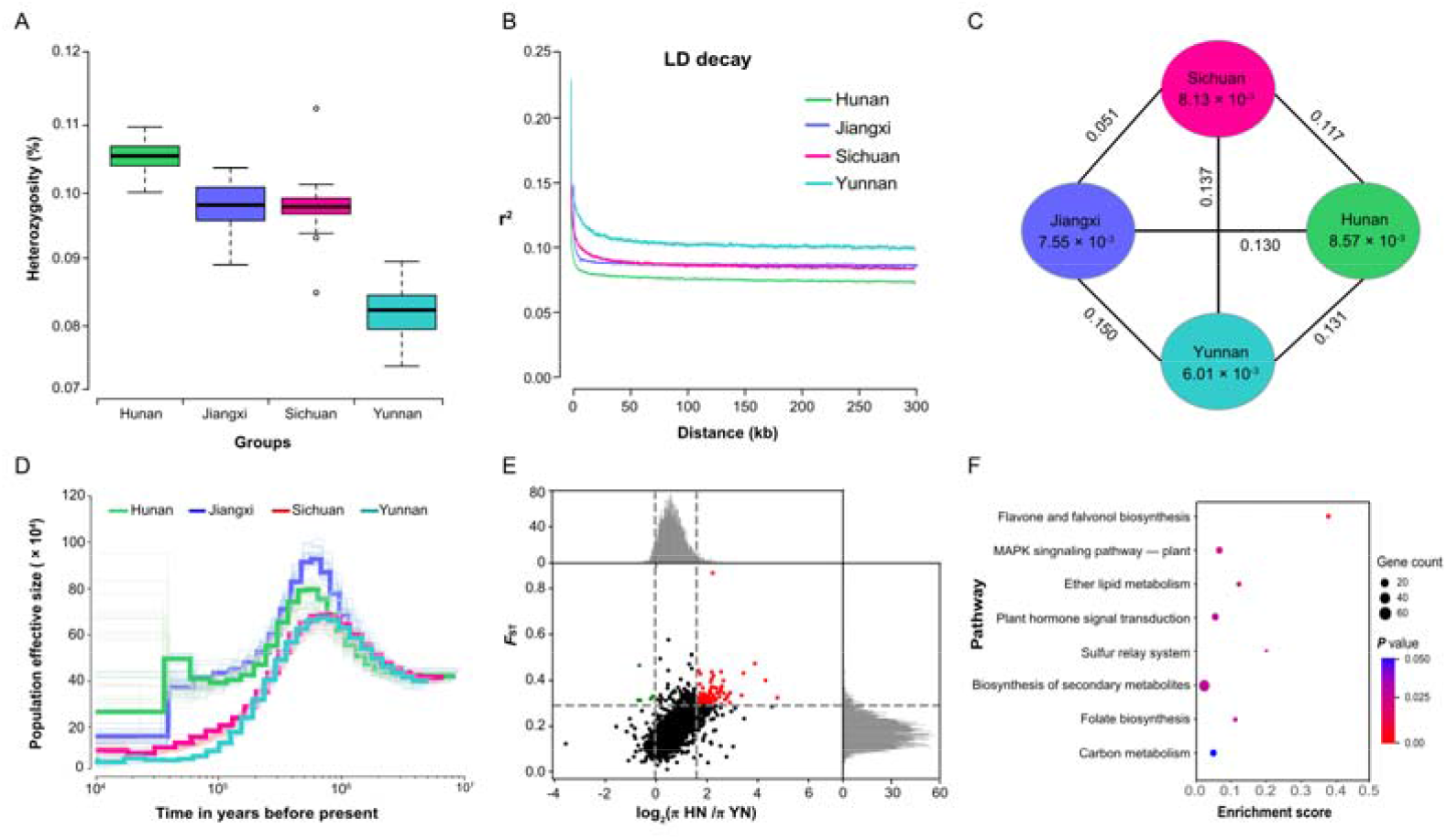
Nucleotide diversity and population divergence of the 101 Shanmei samples. **A**. Genomic heterozygosity of the Hunan, Jiangxi, Sichuan, and Yunnan Shanmei groups. **B**. Decay of LD in four groups of Shanmei. **C**. Nucleotide diversity (π) and population divergence (*F*_ST_) among the four Shanmei groups. Values between pairs indicate population divergence, and values in each circle represent the nucleotide diversity (π) for corresponding group. **D**. Historical effective population size of four Shanmei groups. **E**. Distribution of population differentiation (*F*_ST_) and π ratio (log_2_(π HN/π YN)) between the Hunan and Yunnan groups. *F*_ST_ and π values were calculated across the Shanmei genome using a 50-kb sliding window. **F**. Functional enrichment of genes located at genomic regions under selection in the Yunnan group. LD, linkage disequilibrium;

To reveal the genetic basis of the strong selection in the Yunnan group, the Hunan, Jiangxi, and Sichuan groups were used as controls to determine the candidate genomic regions under selection through genome scanning with a 50-kb sliding window. We found 97 regions that displayed increased levels of differentiation between the Yunnan group and Hunan group (YN_HN) and a significant reduction in nucleotide diversity in Yunnan (*F*_ST_ > 0.29; log2(π HU/π YN) > 1.59; both exceeding the top 5% threshold). Similarly, a total of 94 regions between the Jiangxi group and Yunnan group (JX_YN), and 57 regions between the Sichuan group and Yunnan group (SC_YN) were identified (Figure 4E). In total, we identified 749, 679, and 435 genes in the candidate regions of the HN_YN, JX_YN, and SC_YN comparisons, respectively.

It was found that the flavonoid biosynthesis-related genes were strongly enriched in genes under selection (Figure 4F, Figure S9 and 11). By comparing the nucleotide diversity of genes in the flavonoid biosynthesis pathway (**Figure 5A**), we found that the Yunnan group had the lowest polymorphism (π = 6.1 × 10^−4^), indicating strong selection of these genes in the Yunnan group (Figure 5B). Because the genome-wide diversity of Yunnan group is lower than that of the other groups, we further compared the diversity of flavonoid biosynthesis genes with all genes as the genome background. The results showed that the π values of flavonoid biosynthesis genes were lower than that of the genome background in Yunnan group, which was different to that in the other groups (Figure 5B). Moreover, genes of flavonol synthase (*FLS*) and anthocyanidin synthase (*ANS*) displayed remarkable differences between the Yunnan group and the other groups (Figure 5C). *ANS* is a key component in anthocyanin biosynthesis, which not only is responsible for the coloring of plants [29], but also responds to changes in the external environment [30]. A high abundance of *ANS* enhanced the resistance of bell pepper to low temperature and ultraviolet-B radiation [31]. *FLS* exhibits great potential for regulating plant growth and development, and enhancing plant resistance under abiotic stresses. For example, the increase in *CitFLS* expression promoted fruit ripening during citrus fruit development [32]. *FLS* also can help plants to acclimate to salinity and ultraviolet-B [33]. The purifying selection of *FLS* and *ANS* in the Yunnan group indicated their contribution to the local environmental adaptability of Shanmei in Yunnan.

**Figure 5.**
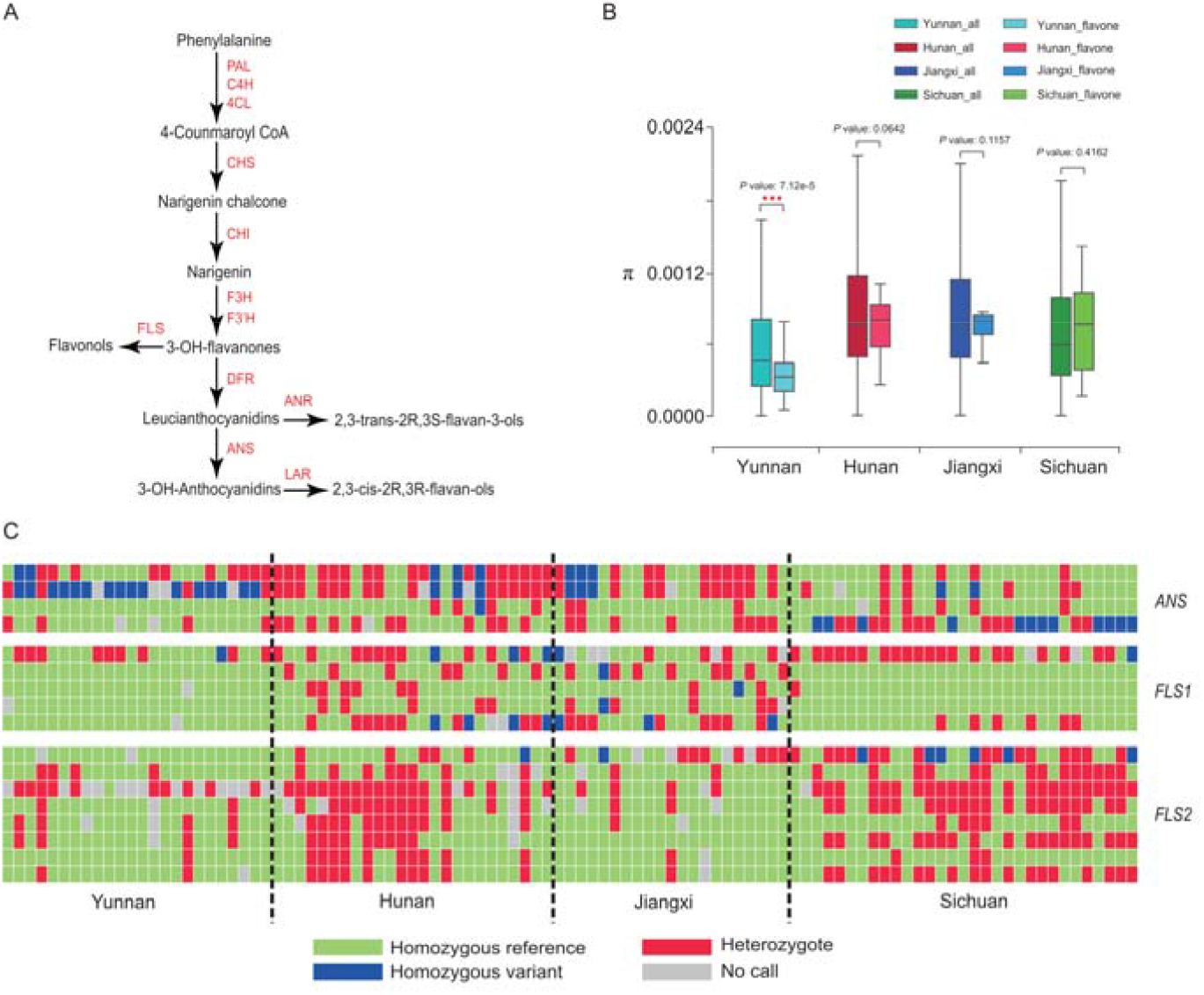
Variations in flavonoid-related genes in the four groups of Shanmei population. **A**. Schematic of the anthocyanin biosynthetic pathway. **B**. Nucleotide diversity (π) comparisons between flavonoid biosynthesis genes and all the genes in the genome of Shanmei for each of the four Shanmei groups. **C**. Genotype variations at non-synonymous SNPs in genes involved in the biosynthesis of flavonoids. *ANS*, Anthocyanidin Synthase; *FLS1*, Flavonol Synthase copy1; *FLS2*, Flavonol Synthase copy 2.

Additionally, some genes related to the mitogen-activated protein kinase (MAPK) signaling pathway and the plant hormone signal transduction pathway were also found to be enriched in these genes under selection. MAPK plays an important role in the plant response to stress. In our study, *MKK2, ANP1*, and *MAPKKK17_18*, key genes in the MAPK signaling pathway [34], were located in regions under selection. Plant hormones are the endogenous messenger molecules that precisely mediate plant growth and development, as well as responses to various biotic and abiotic stresses. Phytohormones play important roles in various biology activities of plants. Genes on the phytohormone signaling pathways were under selection in the Yunnan group, such as genes involved in the abscisic acid (ABA) signaling (*PYL, PP2C*, and *NCED*) and auxin signaling (*IAA, ARF*, and *SAUR*). These results highlighted the importance of the MAPK signaling pathway and plant hormone signal transduction in the environmental adaptability of Shanmei.

## Discussion

Shanmei is a widely distributed wild species that possesses many important characteristics, such as strong adaptability and high medicinal efficacy, thus providing promising genetic materials for breeding. In our study, we assembled a chromosome-scale genome of Shanmei and analyzed its population features. The assembled genome and variome datasets serve as valuable resources for future evolutionary and molecular breeding studies of Shanmei.

The strong environmental adaptability of Shanmei makes it a pioneer plant for reclaiming wasteland primarily due to its high reproduction efficiency and barren tolerance. The assembled genome provides important information on the genetic mechanisms underlying its adaptability. Interspecies comparative genomic analysis revealed that *HSP*s, *RPM1, BIN2*, and *BRI1* underwent significant expansion in gene copy number in Shanmei genome. The *HSP* genes can enhance the heat stress ability of plants [20], while the *RPM1* gene can reduce the damage caused by pathogens [22]. The expression of *BIN2* and *BRI1* that involved in brassinosteroid signal transduction was significantly increased under heat, salt, heavy metal, and drought stress [36, 37]. Furthermore, we found that the copy number of the key genes related to lignin biosynthesis, such as *CAD, CCR, COMT*, and *CCoAOMT*, increased generally in a gradient fashion in herbs, shrubs, and trees. These genes are associated with plant height. For example, the *CAD* and *CCR* mutations displayed a severe dwarfing phenotype [24], and the *COMT* and *CCoAOMT* double mutation resulted in reduced lignin and dwarfing in *Medicago truncatula* [38]. Therefore, we speculated that the increase in gene copy number may lead to an increase in expression dosage, which in turn leads to differences in phenotypes.

The resequencing data further contributed to our understanding of the population divergence and environmental adaptability of Shanmei. Selective sweep analysis focusing on the Yunnan group, which is from the high altitude region and was identified to be under relatively stronger selection compared with the other groups, determined that flavonoid biosynthesis-related genes, as well as genes functioning in plant hormone signal transduction, were enriched in the genomic regions under selection. This indicated that these pathways were crucial to the adaptation of Shanmei to its environment. Generally, flavonoids protect plants against UV, high temperatures, and pathogens [39, 40]. Furthermore, in our study, we found that the Yunnan population exhibited strong selection for genes of *FLS* and *ANS*, which catalyze the biosynthesis of anthocyanins and flavanols, respectively. *FLS* and *ANS* enhance plant resistance to high temperatures and ultraviolet light [41, 42]. Additionally, key genes related to ABA were identified to be under selection in Shanmei, including genes *PYL, PP2C*, and *NCED* that are essential for ABA biosynthesis during salt and drought stress [43, 44]. Taken together, these results suggested that flavonoid biosynthesis and plant hormone signal transduction pathways are important for enhancing the environmental adaptability of Shanmei and serve as potential genetic targets for the further cultivation selection of *Rubus* species.

## Materials and methods

### Materials, sampling, and sequencing

Shanmei seedlings were collected in Jiangxi province of China (115.98°E, 29.68°N) and transplanted into the greenhouse of the Chinese Academy of Agricultural Science. The genomic DNA was isolated from the tender leaves using the DNeasy plant mini kit (Qiagen 69104, Dusseldorf, Germany). The Nanopore library was build according to the manufacturer’s protocol, and genomic sequencing was performed to generate long reads using the Oxford Nanopore PromethION sequencer platform. For Illumina sequencing, a paired-end library was constructed with an insert size of 350 bp and sequenced using the Illumina HiSeq platform, which was used to estimate genomic characteristics and sequence polish. Details of the sequencing are provided in Table S1.

Considering that Shanmei is mainly distributed south to a line from the northeast to the southwest of China (https://www.cvh.ac.cn), samples from four representative regions were collected. They are the Hunan population (114.43°E, 27.29°N, 1000–1300 m) that is located at the central region of South China, the Jiangxi population (115.98°E, 29.68°N, 1100–1300 m) that is located in the east of South China, the Sichuan population (103.22°E, 29.35°N, 1400–1600 m) that is located at the west of South China, and the Yunnan population (104.43°E, 23.15°N, 1700–1900 m) that is located at Southeast China. The Yunnan Shanmei samples distribute in the high altitude region, serving as a subpopulation under specific environmental selection. Two micrograms of DNA per sample was extracted from the fresh leaves using a standard cetyl trimethylammonium bromide (CTAB) extraction protocol. Sequencing libraries were constructed using a Truseq Nano DNA HT Sample Preparation Kit (Illumina USA) following the manufacturer’s instructions. These libraries were sequenced by the Illumina NovaSeq platform, and 150 bp paired-end reads were generated with insert sizes around 350 bp.

### Genome assembly

Jellyfish (version 2.3.0) [45] was used to calculate the k-mer depth distribution with the Illumina short reads, and GenomeScope (version 1.0) [46] was used to estimate the genome size and heterozygosity. Then, NextDenovo (version 2.0, https://github.com/Nextomics/NextDenovo) was used to assemble the Nanopore reads into contigs. The racon (version 1.3.2) [47] and pilon (version 1.2.3) [48] were further used to polish the original contigs with the Nanopore and Illumina reads, each was run for three rounds. Finally, Purge Haplotigs (version 1.2.3) [18] was used to remove heterozygous segments to generate the final contigs. BUSCO was used to assess the completeness of the genome with the embryophyta_odb10 database [49]. Default parameters were used if not specified.

### Hi-C library construction and scaffolding

Fresh leaves from the same Shanmei plant that used for genome sequencing were collected for Hi-C sequencing. The HindIII restriction enzyme was used during the library preparation procedure. The high-quality library was sequenced using the Illumina HiSeq platform. The Hi-C reads were filtered by removing adapter sequences and low-quality reads using Trimmomatic (version 0.39) [50]. The retained Hi-C reads were aligned to the contigs using Juicer (version 1.5, https://github.com/aidenlab/juicer) to obtain the interaction matrix. ALLHIC (version 0.9.8) [51] was used to group, order, and orientate the contigs. Finally, the linking results were manually curated to correct mis-joins and mis-assemblies based on the Hi-C heatmap using JuicerBox (version 1.11.08) [52].

### Repetitive element prediction

LTR_retriever (version 2.7) [53] and RepeatModeler (version 1.0.4) [54] were used to construct the *de novo* repeat libraries. Then, cd-hit software was used to merge the resultant libraries into a non-redundant repeat library (parameters: -c 0.8 -as 0.8 -M 0). Finally, RepeatMasker (version open-4.0.7) [54] was applied to identify and mask the repeat sequences in the Shanmei genome based on the library.

### Protein-coding gene prediction and annotation

An integrated approach was applied to predict the protein-coding genes by merging the results from homology-based searches, mRNA-seq assisted prediction, and *ab initio* prediction. For annotation of homologs, genome sequences of eight species (grape, strawberry, blackberry, apple, peach, pear, apricot, and Chinese rose) were collected from the Genome Database for Rosaceae and were then aligned to Shanmei genome to identify the homologous genes using Exonerate (version 2.4.7) [55]. The *ab initio* gene prediction of Shanmei genome was performed using Genemark (version 4.61_lic) [56] and AUGUSTUS (version 3.3.3) [57]. The RNA-seq data from three tissues (roots, stems, and leaves) were used for transcriptome prediction. Specifically, Hisat2 (version 2.2.1) [58] and Stringtie (version 2.1.4) [59] were used to map RNA-seq reads to the assembled genome and to assemble the alignments into transcripts, respectively. TransDecoder (version 5.5.0, https://github.com/TransDecoder/TransDecoder) was used to identify the potential coding regions in the resultant transcripts. Meanwhile, the RNA-seq reads were *de novo* assembled into transcripts by Trinity (version 2.11.0) [60] using the genome-guided mode, and PASA (version 2.3.1) [61] was used for gene prediction from these transcripts. Finally, EvidenceModeler (version 1.1.1) [62] was used to integrate all gene prediction datasets to generate the final gene set of Shanmei. The predicted protein-coding genes were aligned to the KEGG databases and annotated using KEGG Automatic Annotation Server (KAAS) [63] with an E-value threshold of 1 × 10^−5^.

### Gene expansion and contraction

To identify homologous genes among Shanmei and other plants, the protein sequences of Shanmei were aligned to those of other species (grape, strawberry, blackberry, apple, peach, pear, and Chinese rose) using OrthoFinder (version 2.2.7) [64] with an E-value threshold of 1 × 10^−5^. The protein sequences of single-copy genes were aligned using MUSCLE (version 3.8.31) [65], and the phylogenetic tree was constructed using RAxML (version 8.2.10) [66] with the maximum likelihood algorithm. CAFE (version 4.2.1) [19] was used to identify the expanded and contracted gene families for each species. Default parameters were used if not specified.

### SNP calling and filtering

The paired-end re-sequencing reads were filtered with Trimmomatic (version 0.38) [50]. BWA-MEM (version 0.7.17) [67] was used to align the reads of each sample to the assembled genome. Then, the sequence Alignment (SAM) files were sorted and indexed using samtools (version 1.6) [68]. The GATK (version 1.7.0) [28] genome analysis toolkit was employed to identify variants. In order to obtain high-confidence variants, raw variants were filtered using VCFtools (version 0.1.16) [69]. The filtering criteria were as follows: (1) only SNPs with consensus quality (minQ) ≥ 30 and average SNP depth (minDP) ≥ 10 were retained; (2) the multiallelic sites were filtered out; (3) only SNPs with minor allele frequencies (MAFs) ≥ 0.01 and a minor allele count (mac) ≥ 3 were kept; and (4) SNPs were further filtered based on linkage disequilibrium (LD) with the parameter: --indep-pairwise 100 kb 1 0.5. Finally, 759,241 high-quality SNPs were retained for subsequent analyses. The SNP annotation was performed using ANNOVAR (version 2010Feb15) [70], and SNPs were categorized into intergenic, upstream, downstream region, intron, and exon types based on their relative locations compared with the annotated genes. The SNPs located in coding exons were further separated into synonymous and nonsynonymous SNPs.

### Phylogenetic and population structure

PHYLIP (version 3.696, https://evolution.genetics.washington.edu/phylip.html)was employed to infer the phylogenies of the Shanmei population based on the neighbor-joining algorithm, and MEGA7 (version 7.0) [71] was used to visualize the phylogenetic tree. A PCA of autosomal SNPs was performed using SNPRelate (version 1.28.0) [72]. Structure analysis was performed using ADMIXTURE (version 1.3.0) [73]. The K-values were set from two to seven to estimate the population structure (with the parameters: -geno 0.05 -maf 0.0037 -hwe 0.0001). Finally, the smallest cross-validation (CV) value appeared at *K*□=□4 (Figure S8).

### Inference of the historical population effective size

PSMC (version 0.6.5-r67) [74] was used to estimate the historical effective population size based on the whole-genome resequencing data of the four Shanmei groups. The mutation rate was assumed as μ = 1.9 × 10^−9^ mutations × bp^-1^ × generation^-1^, which was estimated by r8s (version 1.8.1) [75]. One generation was considered as one year. Finally, the script psmc_plot.pl from the PSMC package was used to visualize the results.

### Genome-wide selection signal scanning

To identify genomic regions under selection in Yunnan Shanmei group comparing to the other groups, values of fixation statistic (*F*_ST_) and π were calculated using the VCFtools (version 0.1.16) [69] with a 50 kb nonoverlapping sliding window. Putative selection targets with the top 5% of log_2_ ratios for both π and *F*_ST_ were identified in Yunnan group comparing to each of the other groups. The genes from the genomic regions under selection were analyzed with in-house scripts.

### Identification of key genes in anthocyanin and lignin biosynthesis

The genes involved in anthocyanin and lignin biosynthesis reported in Arabidopsis were collected as references. The BLASTP and SynOrths (version 1.5) [76] tools were used to search the Shanmei genome for homologous genes with an E-value 1 × 10^−20^. Genes supported by both tools were extracted for subsequent analysis using an in-house script. The genes were further confirmed by functional domains prediction in PfamScan (version 1.5, https://www.ebi.ac.uk/Tools/pfa/pfamscan/). The gene *MYB10* was identified based on *RiMYB10* (GenBank ID: 161878916) from red raspberry (*Rubus idaeus*) using mummer (version 4.0.0) [77]. The phylogenetic trees were build using MEGA7 [71] with the neighbor-joining algorithm.

## Data availability

The genome assembly data has been deposited in the Genome Warehouse [78], the resequencing data has been deposited in the Genome Sequence Archive [79], in the National Genomics Data Center [80], China National Center for Bioinformation / Beijing Institute of Genomics, Chinese Academy of Sciences, under the accession numbers GWHBDNY00000000 and CRA003829, respectively. These datasets are publicly accessible at https://bigd.big.ac.cn/gsa.

## CRediT author statement

**Yinqing Yang:** Formal analysis, Investigation, Writing - original draft. **Kang Zhang:** Investigation, Software, Writing - review & editing. **Ya Xiao:** Validation. **Lingkui Zhang:** Methodology. **Yile Huang:** Methodology. **Xing Li:** Investigation. **Shumin Chen:** Investigation. **Yansong Peng:** Resources. **Shuhua Yang:** Conceptualization, Resources. **Yongbo Liu:** Conceptualization, Resources. **Feng Cheng:** Conceptualization, Supervision, Writing - review & editing, Funding acquisition. All authors read and approved the final manuscript.

## Acknowledgments

This work was supported by the grants from the Biodiversity Survey and Assessment Project of the Ministry of Ecology and Environment, China (No. 2019HJ2096001006), and the Science and Technology Innovation Program of the Chinese Academy of Agricultural Sciences, and the Key Laboratory of Biology and Genetic Improvement of Horticultural Crops, Ministry of Agriculture, People’s Republic of China.

## Supplementary Materials

### Supplementary Tables 1-12

**Supplementary Table S1 The statistics of sequencing data used for the Shanmei genome assembly**

**Supplementary Table S2 The length and the number of contigs in each chromosome of Shanmei**

**Supplementary Table S3 The statistics of genes in each prediction process**

**Supplementary Table S4 The statistics of different groups of transposable elements in the genome of Shanmei**

**Supplementary Table S5 The KEGG enrichment analysis of expanded genes in the genome of Shanmei**

**Supplementary Table S6 The KEGG enrichment analysis of contracted genes in the genome of Shanmei**

**Supplementary Table S7 Function annotation of expanded genes in Shanmei**

**Supplementary Table S8 Identification of key genes in anthocyanin biosynthesis**

**Supplementary Table S9 Copy number variation of key genes for lignin biosynthesis in Rosaceae**

**Supplementary Table S10 Resequencing data statistics for the 101 Shanmei samples**

**Supplementary Table S11 Distribution of SNPs in each chromosome of Shanmei**

**Supplementary Table S12 The heterozygosity ratio of each resequenced Shanmei sample**

### Supplementary Figures 1-14

**Supplementary Figure S1 The genome size of Shanmei estimated by GenomeScope**

**Supplementary Figure S2 Whole genome Hi-C contacts of Shanmei**

The black triangles represent the positions of telomere sequences in the seven chromosomes of Shanmei.

**Supplementary Figure S3 Genome synteny analysis of Shanmei, blackberry, and strawberry by MCscanX**

The numbers indicate the chromosome order. The line represents a one-to-one correspondence of homologous regions between genomes of Shanmei and blackberry or strawberry. *Rubus occidentails*, blackberry; *Rubus corchorifolius*, Shanmei; *Fragaria vesca*, strawberry.

**Supplementary Figure S4 Verification of the segmental translocation in chromosome 6 between Shanmei and blackberry using the information of Hi-C contacts**

**A**. The synteny of chromosome 6 between Shanmei and blackberry. The X-axis denotes the chromosome 6 of blackberry (Ro06). The Y-axis denotes the chromosome 6 of Shanmei (Rf06). **B**. The Hi-C heatmap of Shanmei Rf06. **C**. The synteny of chromosome 6 between Shanmei and blackberry after the re-ordering of Shanmei Rf06 following that of blackberry Ro06. **D**. The Hi-C heatmap of Shanmei Rf06 after re-ordering. There are obvious incorrect Hi-C contacts in the re-ordered Rf06 of Shanmei.

**Supplementary Figure S5 Genomic synteny between Shanmei and Fupenzi**

*Rubus corchorifolius*, Shanmei; *Rubus chingii* Hu, Fupenzi.

**Supplementary Figure S6 Verification of the inversions in chromosome 1 between Shanmei and Fupenzi using the information of Hi-C contacts**

**A**. The synteny of chromosome 1 between Shanmei and Fupenzi. The X-axis denotes the chromosome 1 of Fupenzi (LG01). The Y-axis denotes the chromosome 1 of Shanmei (Rf01). **B**. The Hi-C heatmap of Shanmei Rf01. **C**. The synteny of chromosome 1 between Shanmei and Fupenzi after the re-ordering of Shanmei Rf01 following that of Fupenzi LG01. **D**. The Hi-C heatmap of Shanmei Rf01 after re-ordering.

**Supplementary Figure S7 Verification of the inversion in chromosome 4 between Shanmei and Fupenzi using information of Hi-C contacts**

**A**. The synteny of chromosome 4 between Shanmei and Fupenzi. The X-axis denotes the chromosome 4 of Fupenzi (LG04). The Y-axis denotes the chromosome 4 of Shanmei (Rf04). **B**. The Hi-C heatmap of Shanmei Rf04. **C**. The synteny of chromosome 4 between Shanmei and Fupenzi after the re-ordering of Shanmei Rf04 following that of Fupenzi LG04. **D**. The Hi-C heatmap of Shanmei Rf04 after re-ordering.

**Supplementary Figure S8 KEGG enrichment analysis of genes sets in Shanmei**

**A**. KEGG enrichment for expanded genes. **B**. KEGG enrichment for contracted genes.

**Supplementary Figure S9 Variations on copy number and expression of key genes involved in lignin biosynthesis in Rosaceae**

**A**. The genes reported in the biosynthesis pathway of lignin. **B**. Heatmap of the copy number of key genes for lignin biosynthesis in Rosaceae. **C**. Phylogenetic tree of the COMT gene family. **D**. The genes’ expression level (TPM: transcripts per million mapped reads) measured by mRNA-seq data of stem organ from three representative species of macrophanerophytes, shrub, and herb. Rf, Shanmei; Fv, strawberry; Pp, peach; Pyc, pear; At, Arabidopsis.

**Supplementary Figure S10 Characterization of MYB10 in Rosaceae species**

**A**. Protein sequence alignment of the MYB10 transcription factors, showing only the part of R2 and R3 domains. Conserved tryptophan residues in the R2 and R3 domains were marked with asterisks (*). The characteristic amino acids in the dicot anthocyanin-promoting MYB transcription factors were highlighted by red boxes. **B**. The RuMYB10 protein 3D structure. The arrow pointed at the Asparagine (N). *Rubus occidentalis*, blackberry; *Rubus corchorifolius*, Shanmei; *Rubus idaeus*, red raspberry; *Fragaria vesca*, strawberry; *Prunus dulcis*, Almod; *Prunus persica*, peach; *Pyrus avium*, Sweet cherry; *Pyrus communis*, pear; *Pyrus pyrifolia*, sand pear; *Malus domestica*, apple.

**Supplementary Figure S11 Standard error estimation of Shanmei population admixture analysis**

**Supplementary Figure S12 KEGG enrichment analysis of genes located at genomic regions under selection in Shanmei Yunnan group comparing to that of Hunan group**

**Supplementary Figure S13 KEGG enrichment analysis of genes located at genomic regions under selection in Shanmei Yunnan group comparing to that of Jiangxi group**

**Supplementary Figure S14 KEGG enrichment analysis of genes located at genomic regions under selection in Shanmei Yunnan group comparing to that of Sichuan group**

