## Supplementary Figures for "Genome Assembly and Population Resequencing Reveal the Geographical Divergence of ‘Shanmei’ (*Rubus corchorifolius*)"

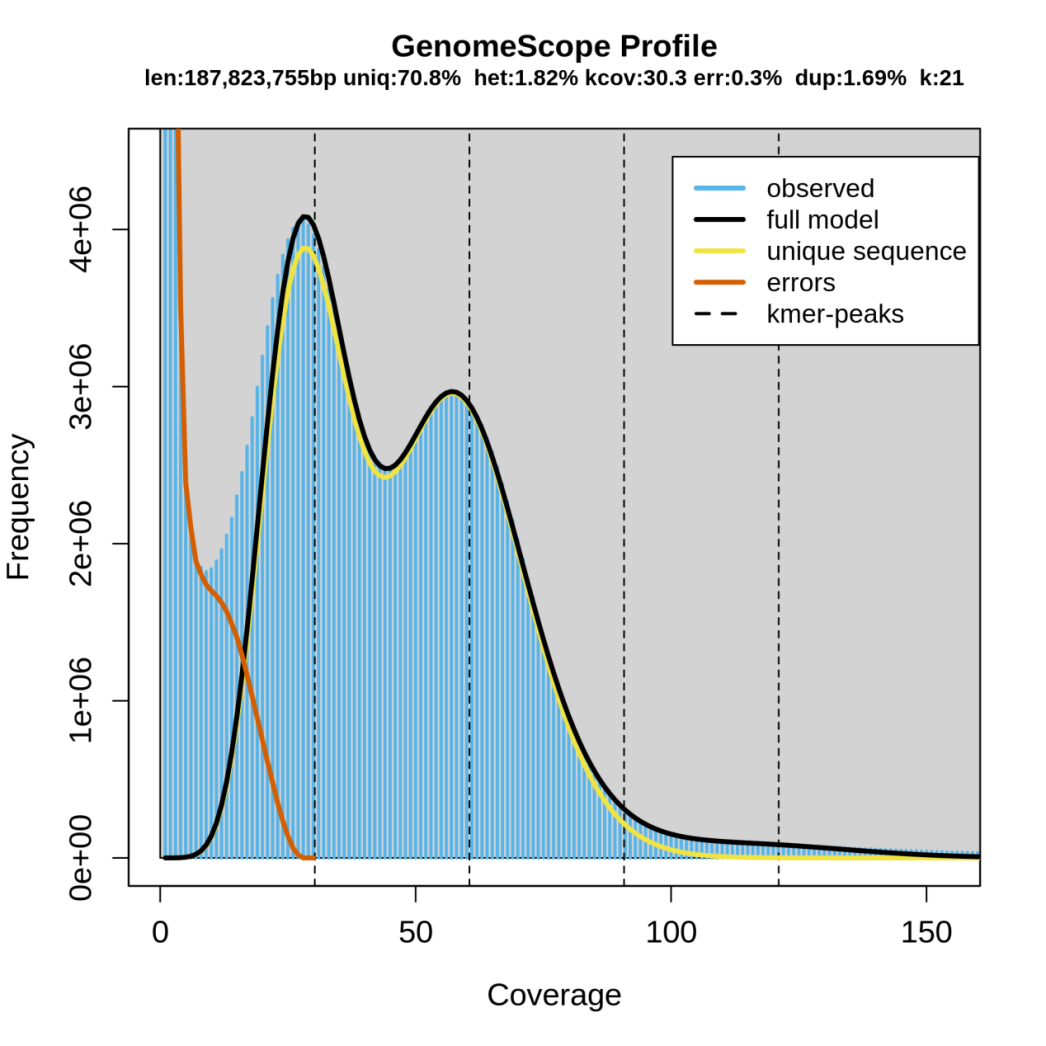

Supplementary Figure 1 **The genome size of Shanmei estimated by GenomeScope**

The predicted genome size was approximately 187.82 Mb, with a heterozygosity ratio of 1.82%.

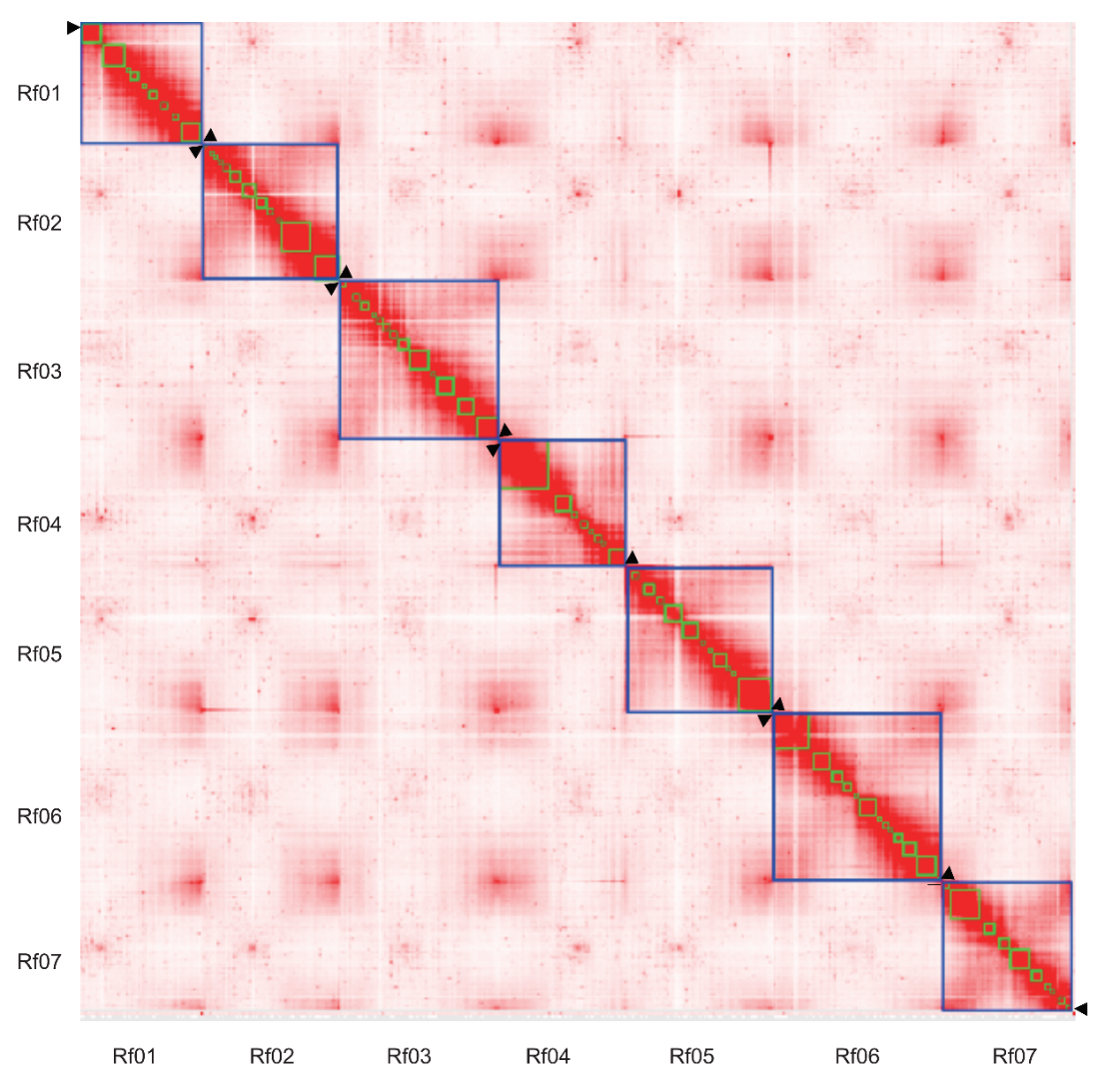

Supplementary Figure 2 **Whole genome Hi-C contacts of Shanmei**

The black triangles represent the positions of telomere sequences in the seven chromosomes of Shanmei.

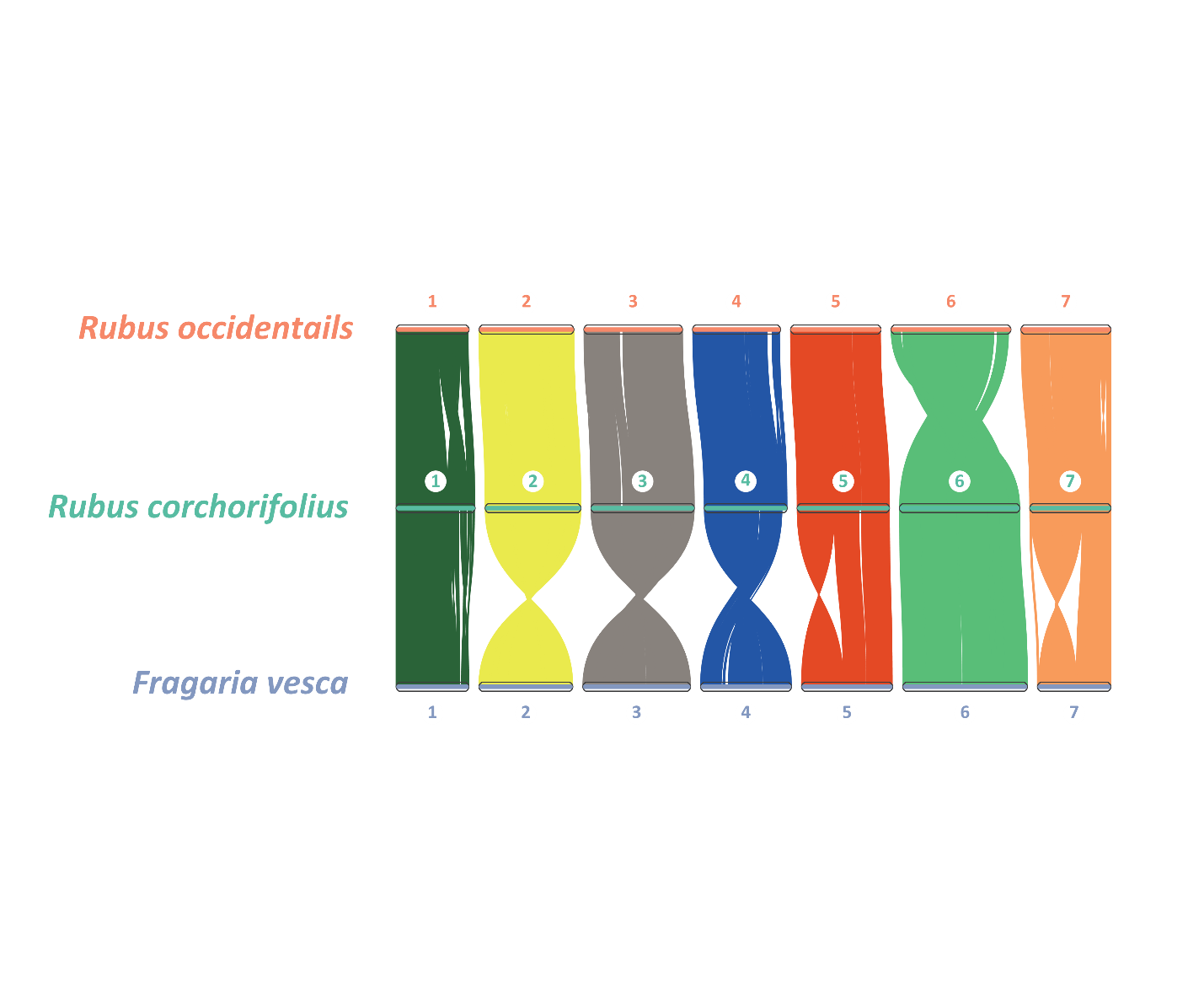

Supplementary Figure 3 **Genome synteny analysis of** **Shanmei, blackberry, and strawberry** **by MCscanX**

The numbers indicate the chromosome order. The line represents a one-to-one correspondence of homologous regions between genomes of Shanmei and blackberry or strawberry. R*ubus occidentails*: blackberry; *Rubus corchorifolius*: Shanmei; *Fragaria vesca*: strawberry.

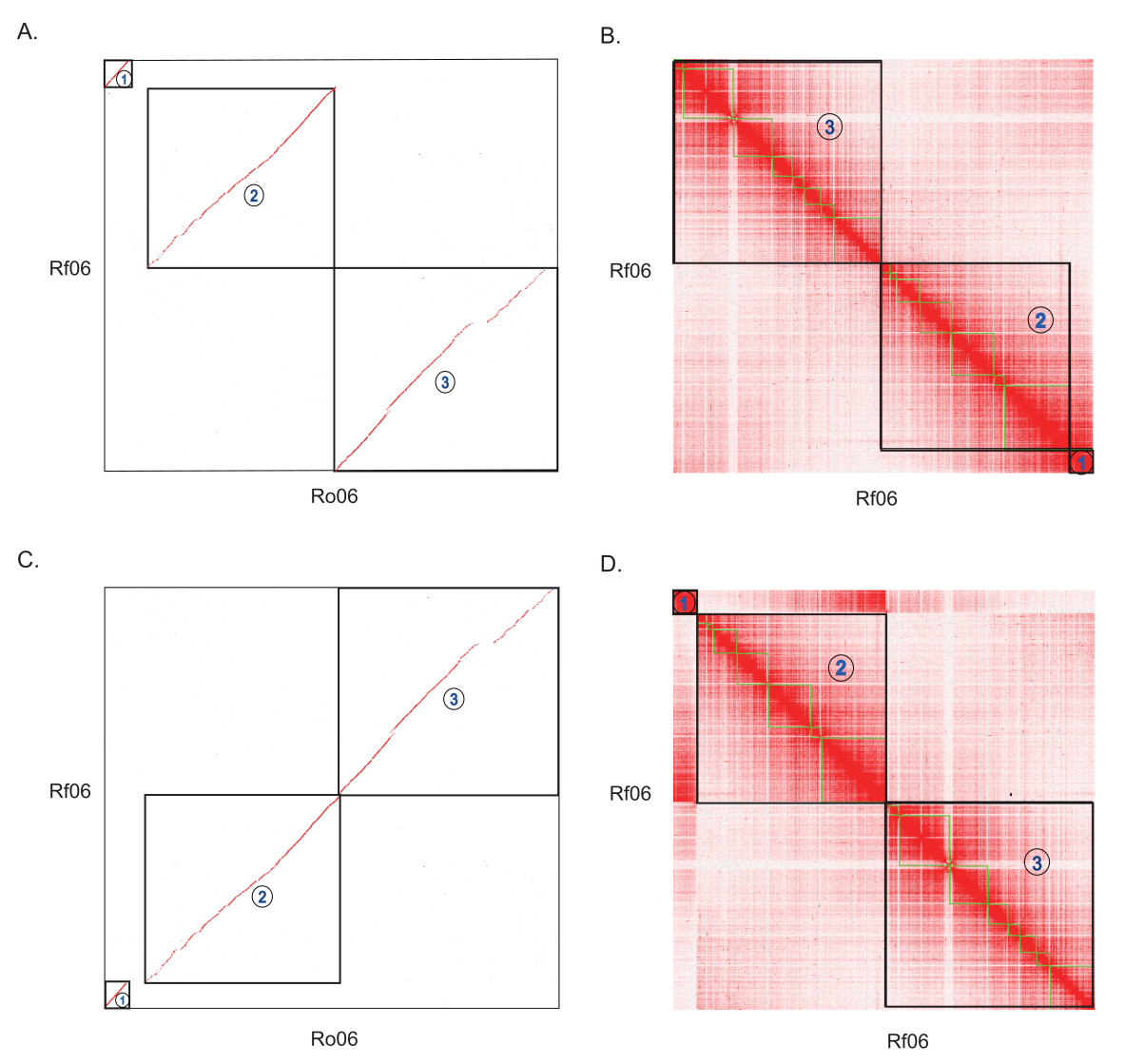

**Supplementary Figure 4 Verification of the segmental translocation in chromosome 6 between Shanmei and blackberry using the information of Hi-C contacts**

**A**. The synteny of chromosome 6 between Shanmei and blackberry. The X-axis denotes the chromosome 6 of blackberry (Ro06). The Y-axis denotes the chromosome 6 of Shanmei (Rf06). **B**. The Hi-C heatmap of Shanmei Rf06. **C**. The synteny of chromosome 6 between Shanmei and blackberry after the re-ordering of Shanmei Rf06 following that of blackberry Ro06. **D**. The Hi-C heatmap of Shanmei Rf06 after re-ordering. There are obvious incorrect Hi-C contacts in the re-ordered Rf06 of Shanmei.

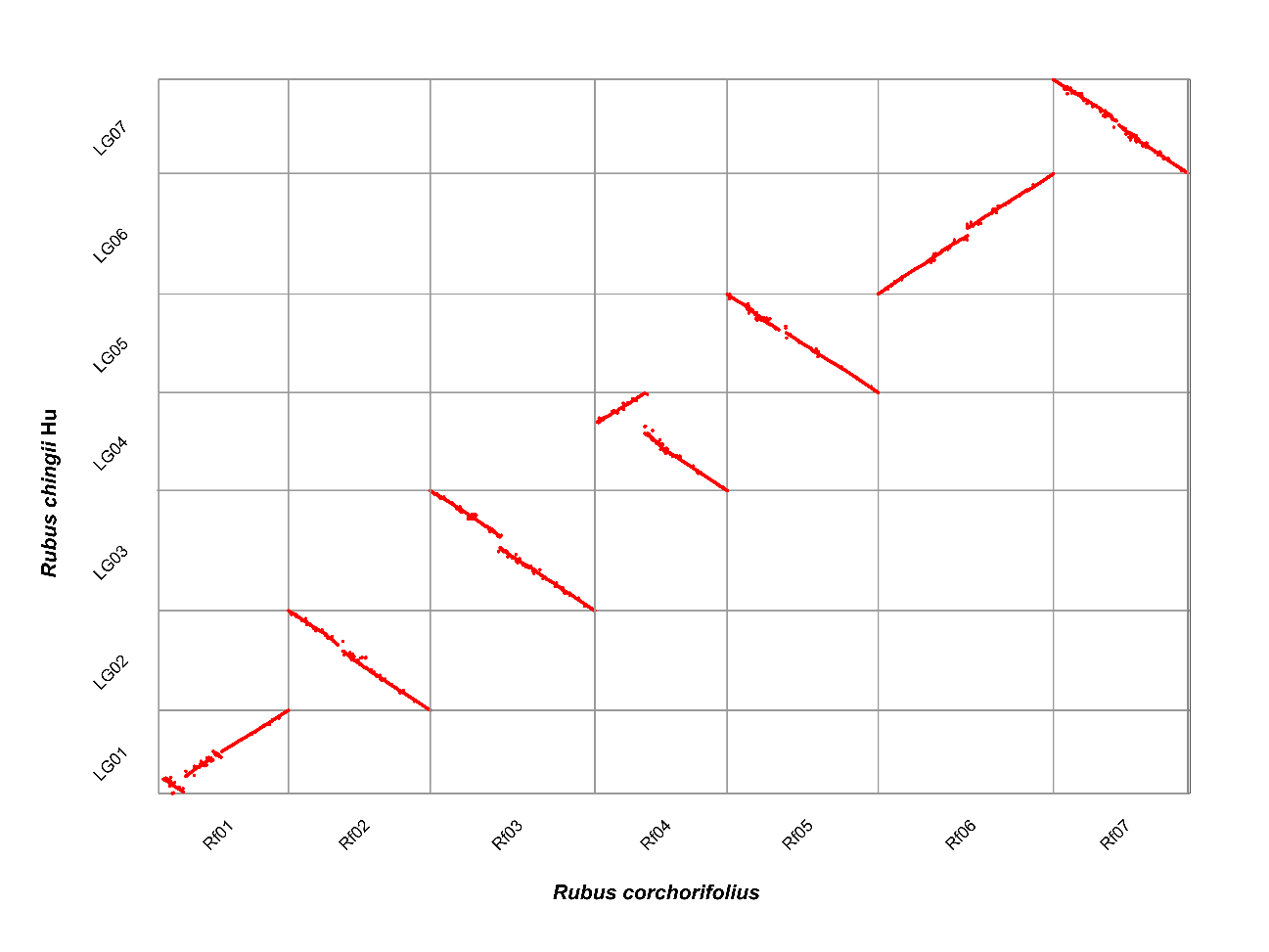

Supplementary Figure 5 **Genomic synteny between Shanmei and Fupenzi**

*Rubus corchorifolius*: Shanmei; *Rubus chingii* Hu: Fupenzi.

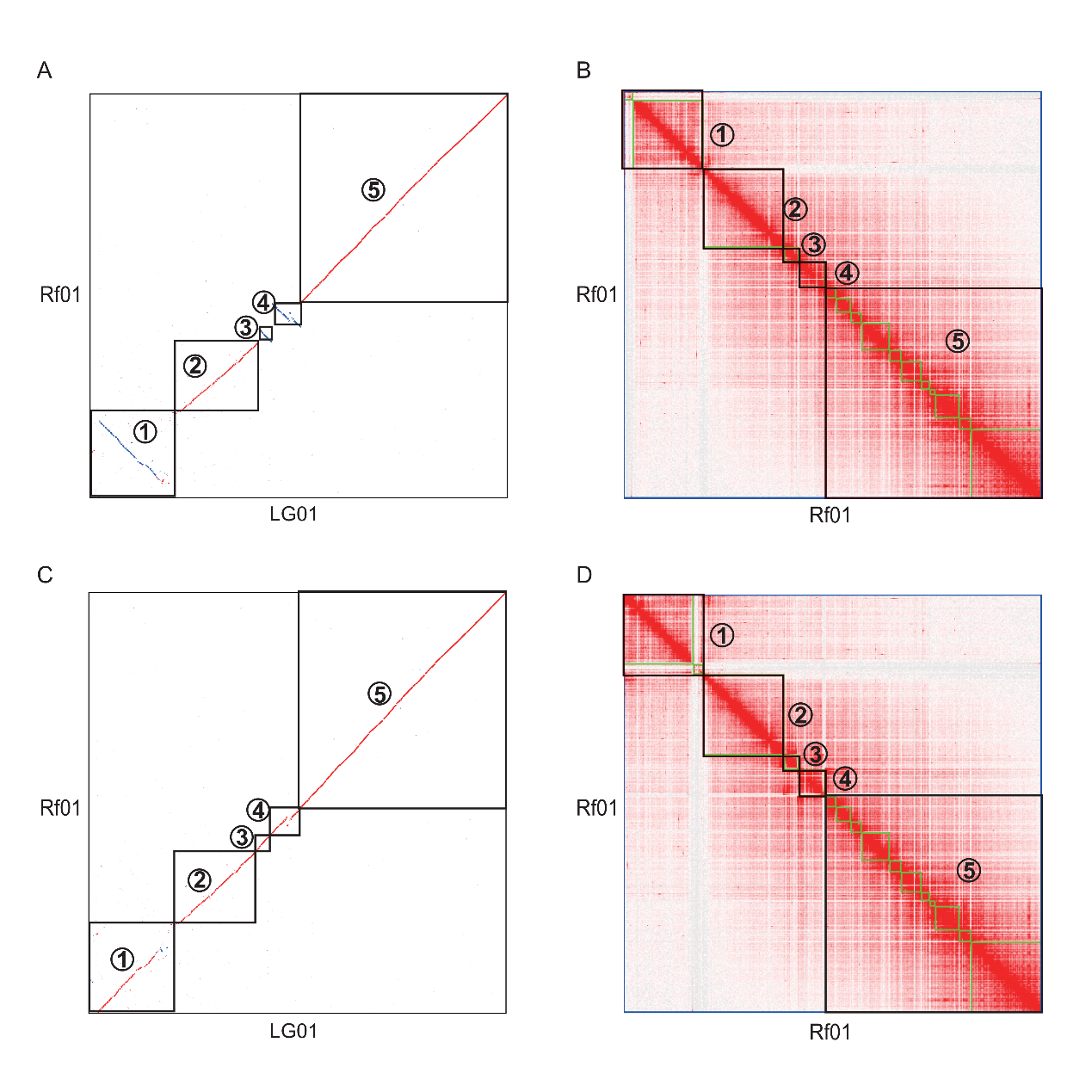

**Supplementary Figure 6 Verification of the inversions in chromosome 1 between Shanmei and Fupenzi using the information of Hi-C contacts**

**A**. The synteny of chromosome 1 between Shanmei and Fupenzi. The X-axis denotes the chromosome 1 of Fupenzi (LG01). The Y-axis denotes the chromosome 1 of Shanmei (Rf01). **B**. The Hi-C heatmap of Shanmei Rf01. **C**. The synteny of chromosome 1 between Shanmei and Fupenzi after the re-ordering of Shanmei Rf01 following that of Fupenzi LG01. **D**. The Hi-C heatmap of Shanmei Rf01 after re-ordering.

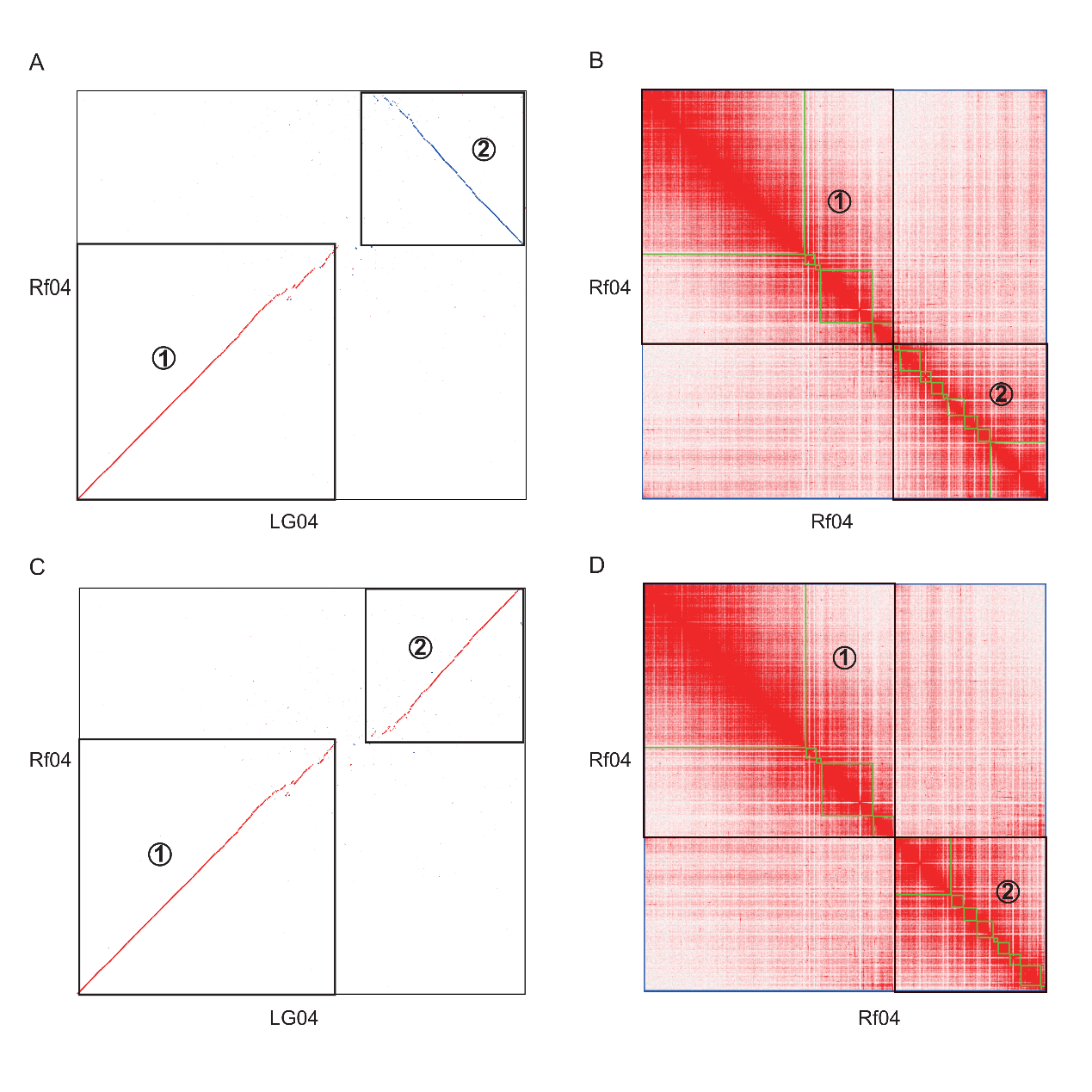

**Supplementary Figure 7 Verification of the inversion in chromosome 4 between Shanmei and Fupenzi using information of Hi-C contacts**

**A**. The synteny of chromosome 4 between Shanmei and Fupenzi. The X-axis denotes the chromosome 4 of Fupenzi (LG04). The Y-axis denotes the chromosome 4 of Shanmei (Rf04). **B**. The Hi-C heatmap of Shanmei Rf04. **C**. The synteny of chromosome 4 between Shanmei and Fupenzi after the re-ordering of Shanmei Rf04 following that of Fupenzi LG04. **D**. The Hi-C heatmap of Shanmei Rf04 after re-ordering.

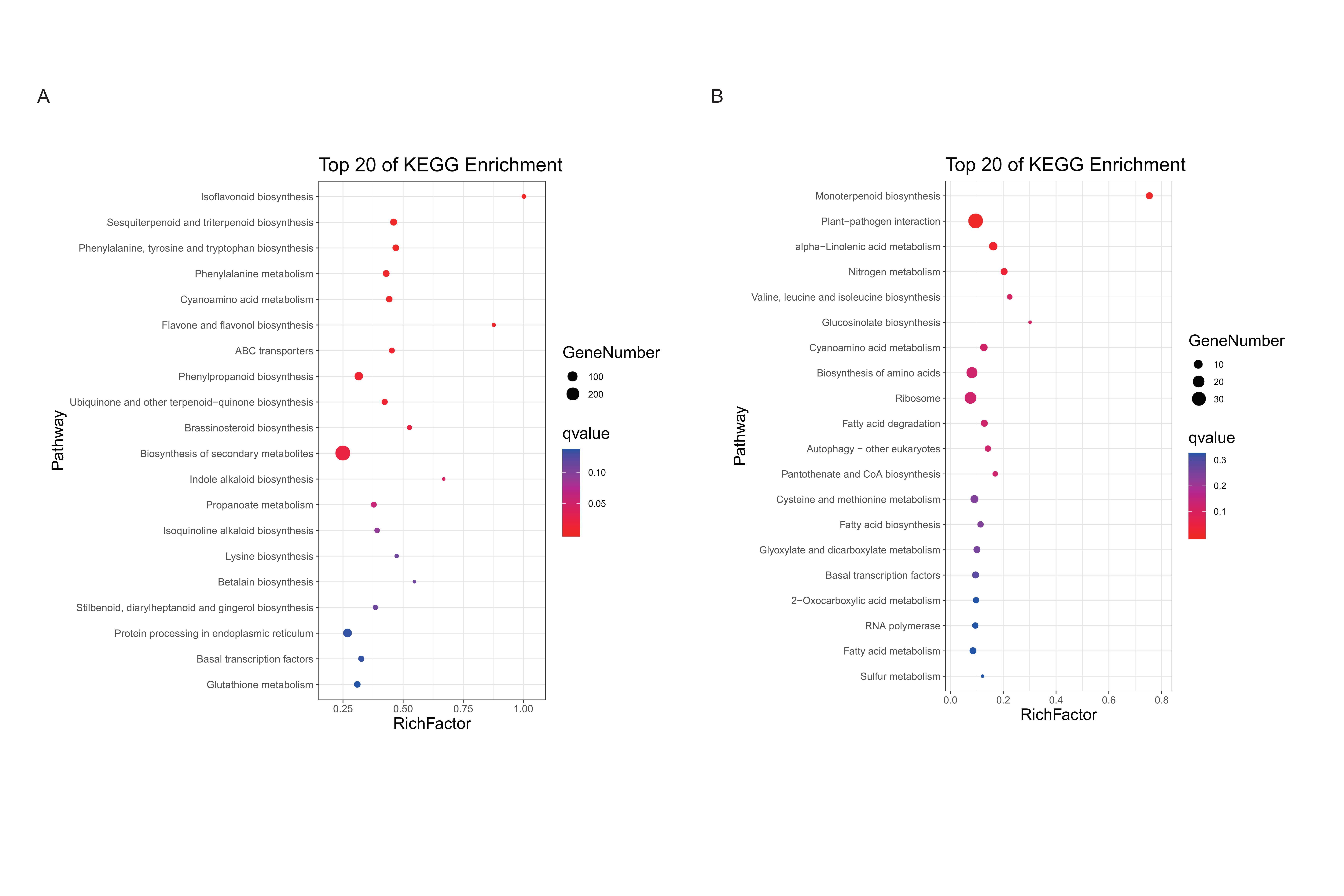

**Supplementary Figure 8 KEGG enrichment analysis of genes sets in Shanmei**

**A**. KEGG enrichment for expanded genes. **B**. KEGG enrichment for contracted genes.

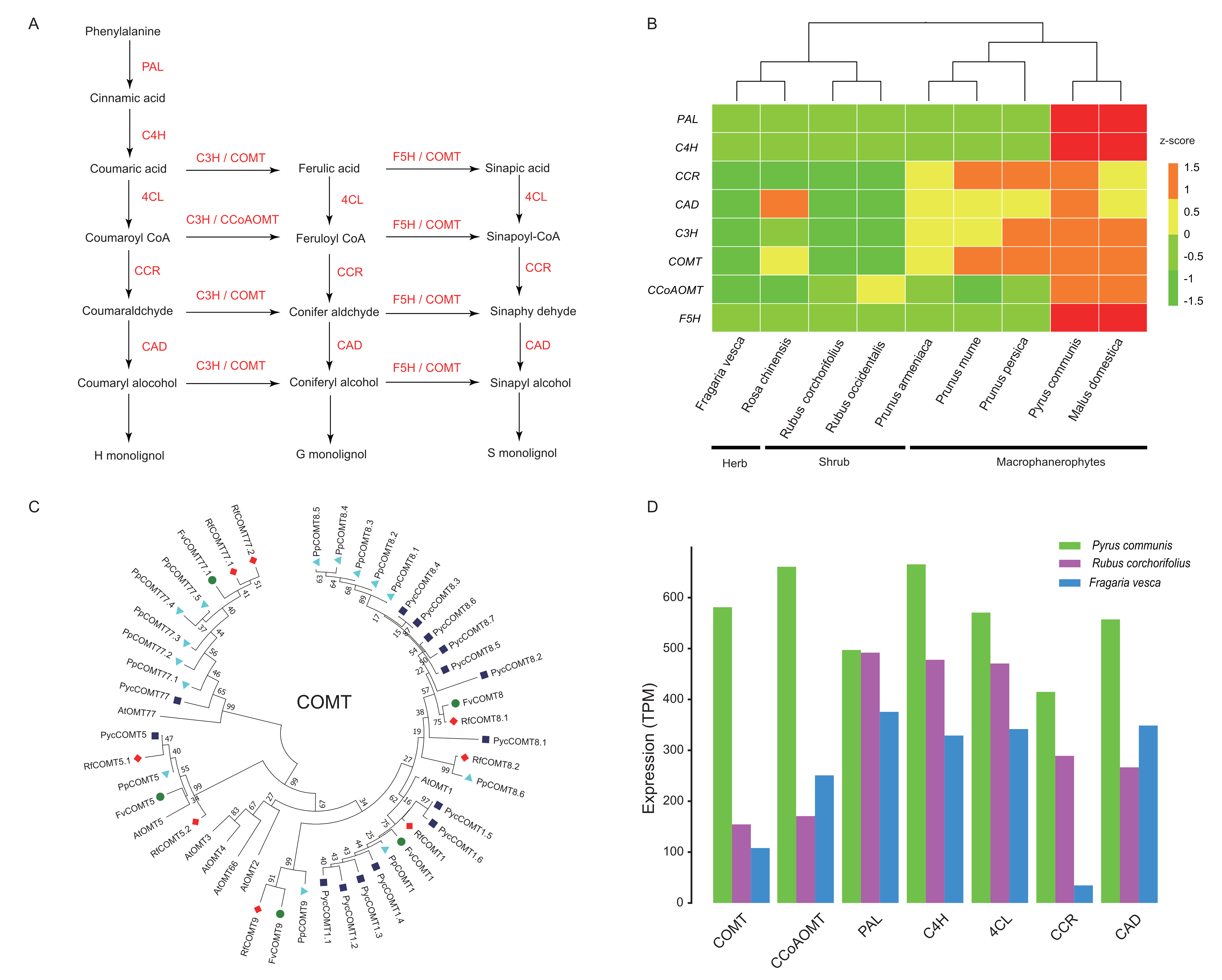

**Supplementary Figure 9**  **Variations on copy number and expression of key genes involved in lignin biosynthesis in Rosaceae**

**A**. The genes reported in the biosynthesis pathway of lignin. **B**. Heatmap of the copy number of key genes for lignin biosynthesis in Rosaceae. **C**. Phylogenetic tree of the COMT gene family. **D**. The genes’ expression level (TPM: transcripts per million mapped reads) measured by mRNA-seq data of stem organ from three representative species of macrophanerophytes, shrub, and herb. Rf: Shanmei; Fv: strawberry; Pp: peach; Pyc: pear; At: Arabidopsis.

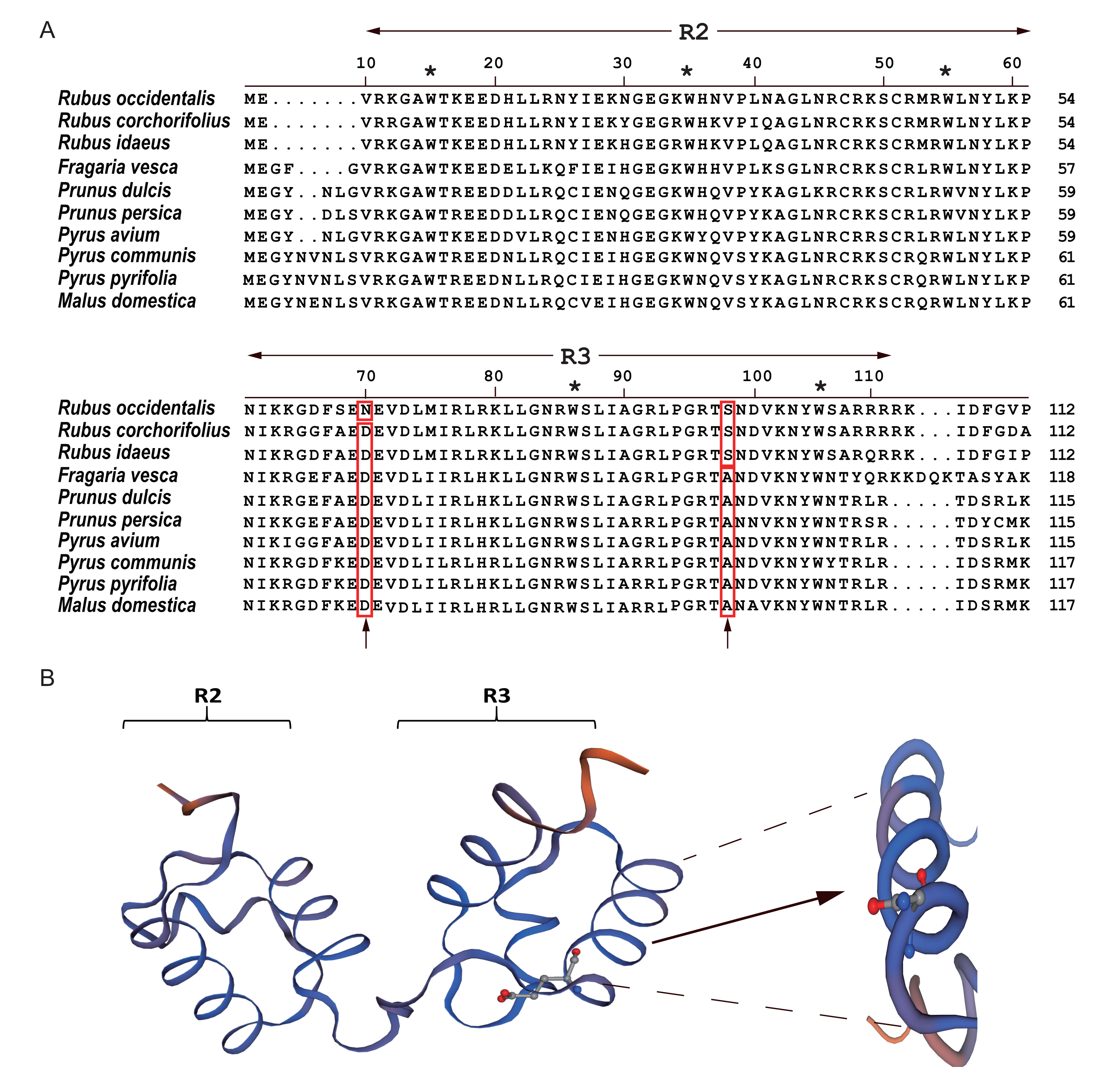

**Supplementary Figure 10 Characterization of MYB10 in Rosaceae species**

**A**. Protein sequence alignment of the MYB10 transcription factors, showing only the part of R2 and R3 domains. Conserved [tryptophan](https://www.sciencedirect.com/topics/agricultural-and-biological-sciences/tryptophan" \o "Learn more about Tryptophan from ScienceDirect's AI-generated Topic Pages) residues in the R2 and R3 domains were marked with asterisks (*). The characteristic amino acids in the [dicot](https://www.sciencedirect.com/topics/agricultural-and-biological-sciences/magnoliopsida" \o "Learn more about Magnoliopsida from ScienceDirect's AI-generated Topic Pages) anthocyanin-promoting MYB transcription factors were highlighted by red boxes. **B**. The RuMYB10 protein 3D structure. The arrow pointed at the Asparagine (N). *Rubus occidentalis*: blackberry; *Rubus corchorifolius*: Shanmei; *Rubus idaeus*: red raspberry; *Fragaria vesca*: strawberry; *Prunus dulcis*: Almod; *Prunus persica*: peach; *Pyrus avium*: Sweet cherry; *Pyrus communis*: pear; *Pyrus pyrifolia*: sand pear; *Malus domestica*: apple.

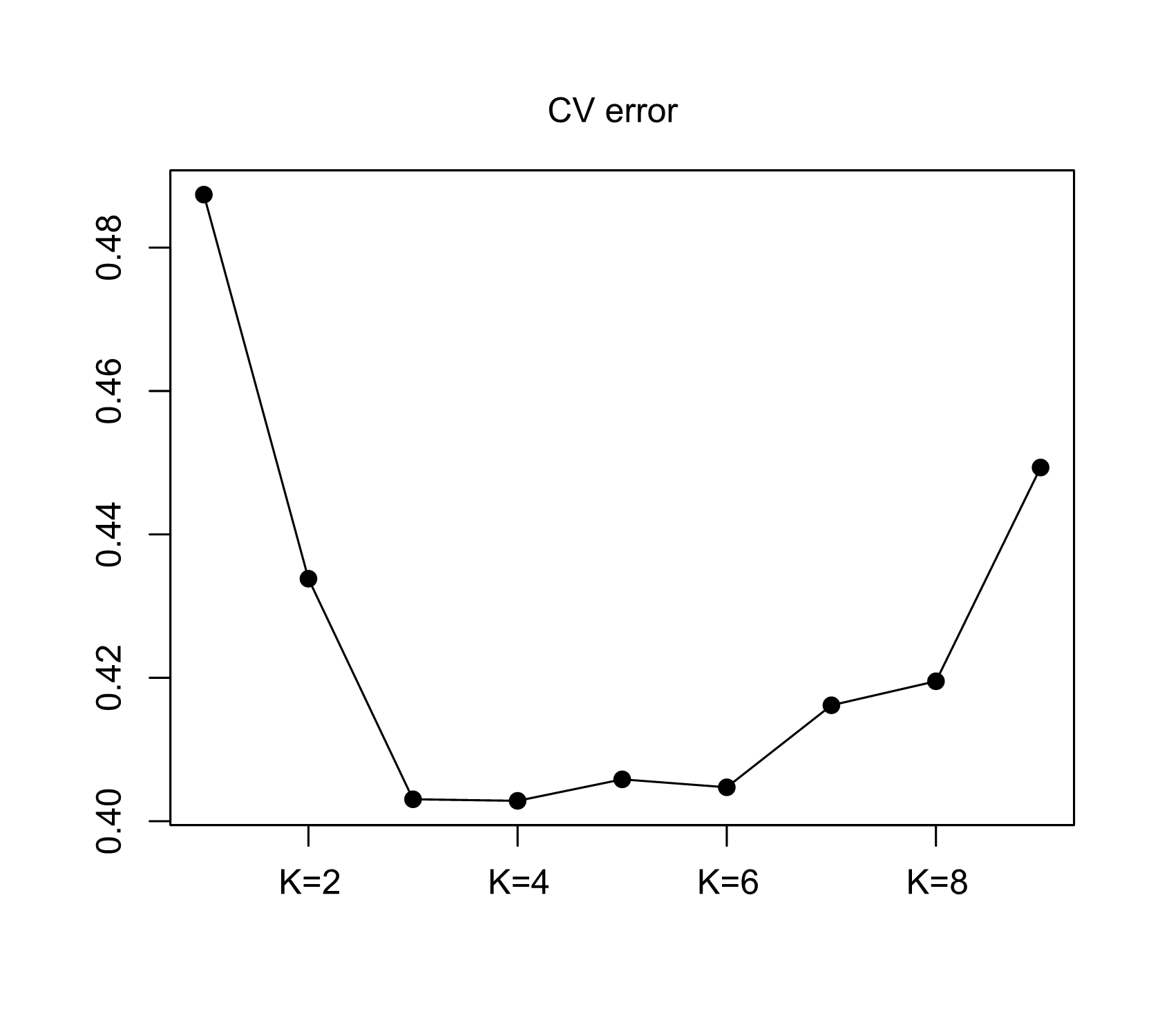

Supplementary Figure 11 **Standard error estimation of Shanmei population admixture analysis**

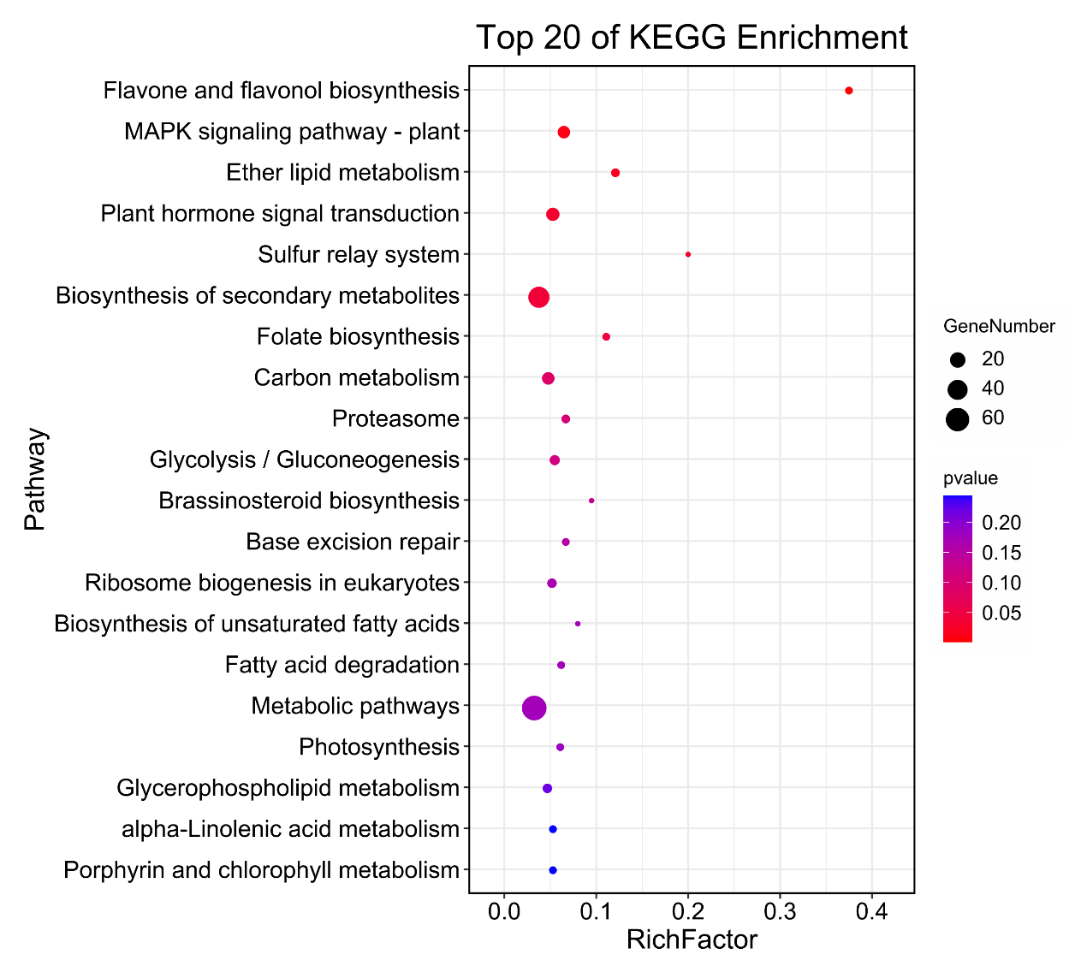

Supplementary Figure 12 **KEGG enrichment analysis of genes located at genomic regions under selection in Shanmei Yunnan group comparing to that of Hunan group**

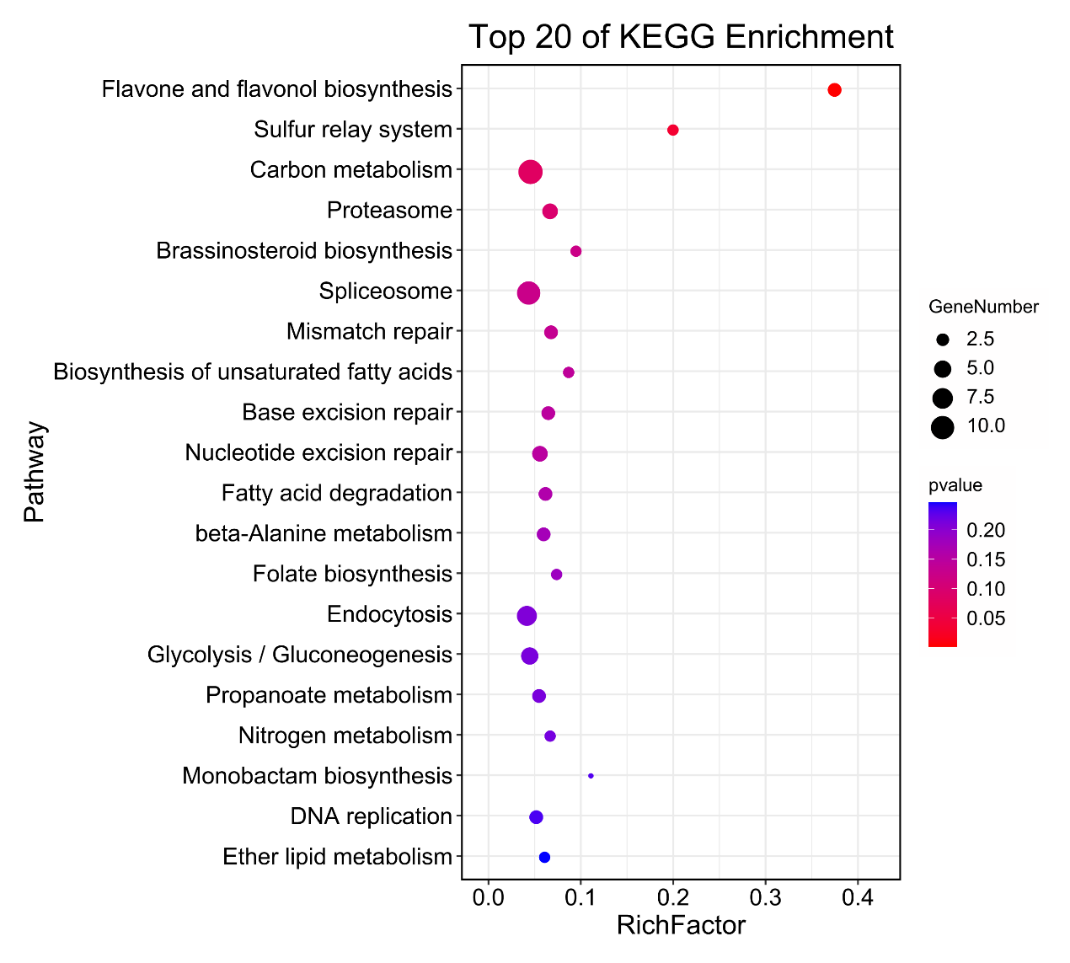

Supplementary Figure 13 **KEGG enrichment analysis of genes located at genomic regions under selection in Shanmei Yunnan group comparing to that of Jiangxi group**

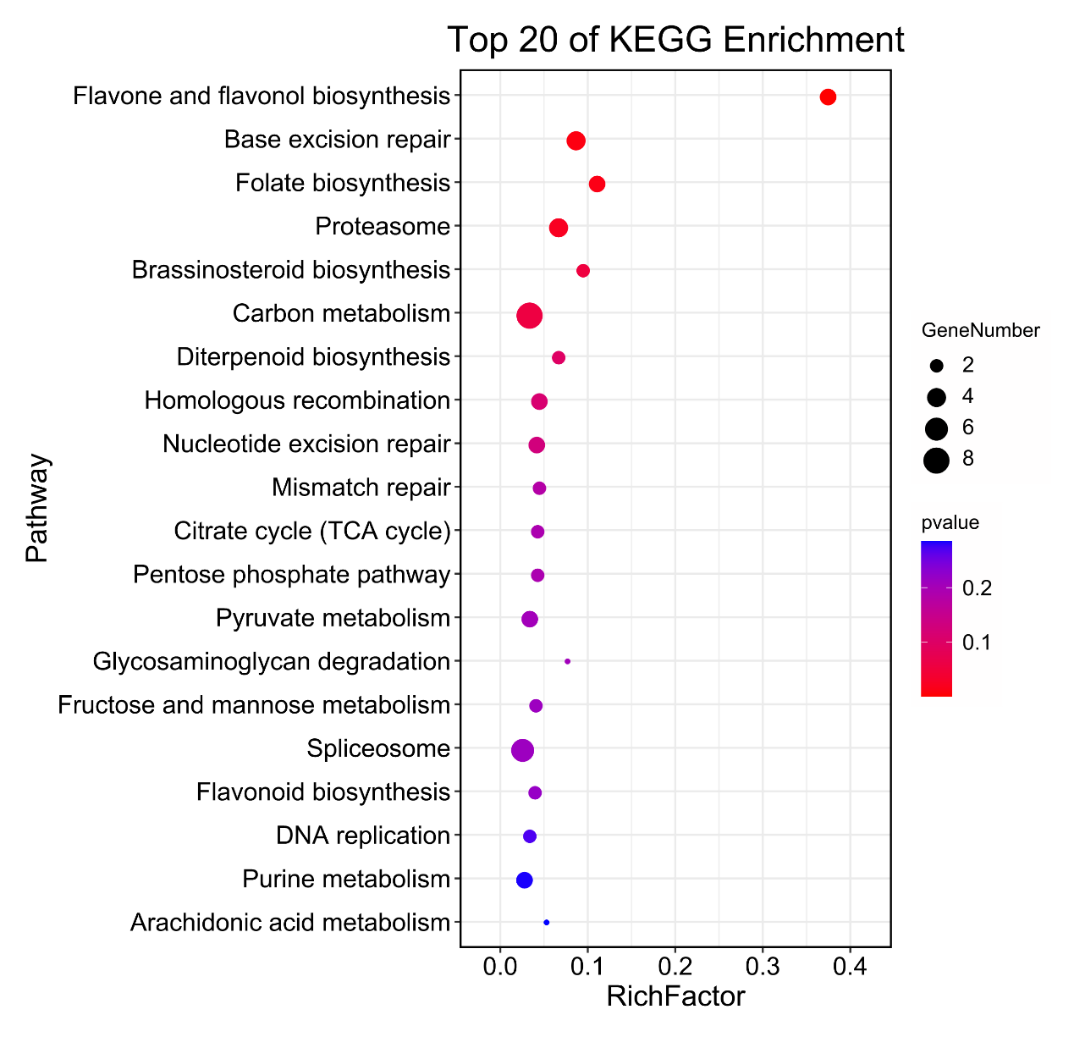

Supplementary Figure 14 **KEGG enrichment analysis of genes located at genomic regions under selection in Shanmei Yunnan group comparing to that of Sichuan group**
